# Cerberus: bidirectional state space blocks improve accuracy and efficiency of regulatory sequence models

**DOI:** 10.64898/2026.10.02.756377

**Authors:** David R. Kelley, Han Yuan, Xingfan Huang, Johannes Linder

**Author notes:** Contributing authors.

## Abstract

Sequence-to-function models predict regulatory activity and variant effects from DNA sequence alone, and leading models such as Borzoi compute long-range interactions with self-attention. The reference genome caps unique training sequences, and new assays add labels, not sequences. This constraint favors architectures that learn more from limited sequence diversity. We compared long-range blocks with the rest of the architecture and the training data held fixed, replacing Borzoi’s transformer blocks with Hydra, a bidirectional state space block built on Mamba-2. Hydra blocks ran 5.7-fold faster on 8,192-token inputs, and models built on them predicted binned coverage on held-out sequences and classified fine-mapped eQTLs more accurately. Hydra’s linear cost in sequence length let us run the blocks at 32 bp resolution and remove the U-net decoder, leaving a convolution-SSM model smaller and more accurate than the one it replaced. Scaling this architecture up on an augmented track collection produced Cerberus, an ensemble of eight models over 786 kb of input. We evaluated it on GTEx v11 fine-mapped eQTL, sQTL, and paQTL benchmarks with negatives matched on allele frequency, gene expression, and phenotype-specific positional annotations to reduce confounding by these properties. Cerberus is more accurate than Borzoi on all three phenotypes, 0.692 versus 0.668 mean eQTL AUPRC across 48 tissues, and estimates eQTL effect sizes better, 0.379 versus 0.321 Spearman ***ρ***. Against AlphaGenome, it trades leads in eQTL classification, ahead on variants 3 to 100 kb from the transcription start site and behind in coding sequence and within 3 kb. Interpretation of the trained blocks shows interaction range growing with block depth, from a 0.9 kb median half-life in the first block to 79 kb in the seventh, and the deepest heads anchoring on promoters, enhancers, and CTCF peaks. We release the models, the training data, the training and evaluation code, and the benchmark sets, supporting variant scoring and transfer learning.

## 1 Background

Long-range sequence-to-function models such as Borzoi [1] and AlphaGenome [2] predict the effects of genetic variants on gene expression, splicing, and polyadenylation from DNA sequence alone. These predictions complement association evidence across the allele-frequency spectrum, informing fine mapping [3, 4], noncoding variant prioritization in rare disease, and variant selection for aggregate association tests [5, 6].

However, the computational resources these models require during both training and inference constrain them in three ways. First, costly training runs undersample the hyperparameter space, so published configurations reflect the search that was affordable, not an optimum. Second, training requires expensive, in-demand hardware, placing it out of reach for many research groups. Third, slow inference bounds how many variants can be scored, a bottleneck for biobank-scale application.

Self-attention drives much of that cost, because its compute grows quadratically with sequence length. To limit it, models such as Borzoi perform long-range computation on coarsely pooled sequence, pooling with a convolution encoder and recovering fine-resolution predictions with a U-net decoder. Both components are themselves expensive. Sub-quadratic sequence mixers now match attention on long-context language tasks [7, 8], and several have been adapted to biological sequence architectures [9–12].

We replaced Borzoi’s transformer blocks with bidirectional state space blocks, specifically Hydra, built on Mamba-2 [7, 13, 14]. A reduced collection of output tracks made the hyperparameter search affordable. Across four training replicates, the substitution improved accuracy in predicting 32 bp coverage on held-out sequences and in classifying fine-mapped eQTLs. Hydra’s linear cost in sequence length then permitted a simpler architecture, operating on finer-resolution sequences without a U-net decoder and with fewer parameters than the model it replaced. To scale this architecture, we augmented the Borzoi training data with posttranscriptional and single-cell atlas-derived tracks and trained Cerberus, an ensemble of eight regulatory foundation models. Basic variant properties separate fine-mapped positives from unselected variants without predicting their regulatory effects. We therefore built new GTEx v11 fine-mapped eQTL, sQTL, and paQTL benchmarks matching every positive to a negative on region, distance, allele frequency, and expression. Cerberus is more accurate than Borzoi on all three phenotypes and trades leads with AlphaGenome across eQTL variant classes. We release the models, the training data, the training and evaluation code, and the benchmark sets open-source, to support variant scoring, transfer learning [15], and continued model development.

## 2 Results

### 2.1 Long-range block comparison and architecture search

To compare alternative long-range blocks, we built a dataset representative of large-scale regulatory sequence modeling with the output tracks downsampled to speed up training. We aligned other major architecture components to Borzoi—524 kb DNA input sequence processed by a convolution encoder to 128 bp resolution, long-range computation, U-net decoder to 32 bp experimental track prediction (Fig. 1a). To train fast enough to explore many alternatives, we capped models at 768 channels (half of Borzoi and AlphaGenome) and six long-range blocks. For each alternative, we tuned learning rate, momentum, and weight decay by manual iterative search, and trained under both a cosine decay schedule and a schedule-free optimizer. We selected a model configuration within each family using average validation set Pearson correlation (r) at bin-level across tracks. Full evaluation considered test set r, specificity-focused r after quantile normalization and mean subtraction within data collections, analogous metrics after gene exon aggregation, and metrics describing genetic variant predictions for GTEx v11 fine-mapped eQTLs and carefully matched negatives (Methods).

**Fig. 1.**
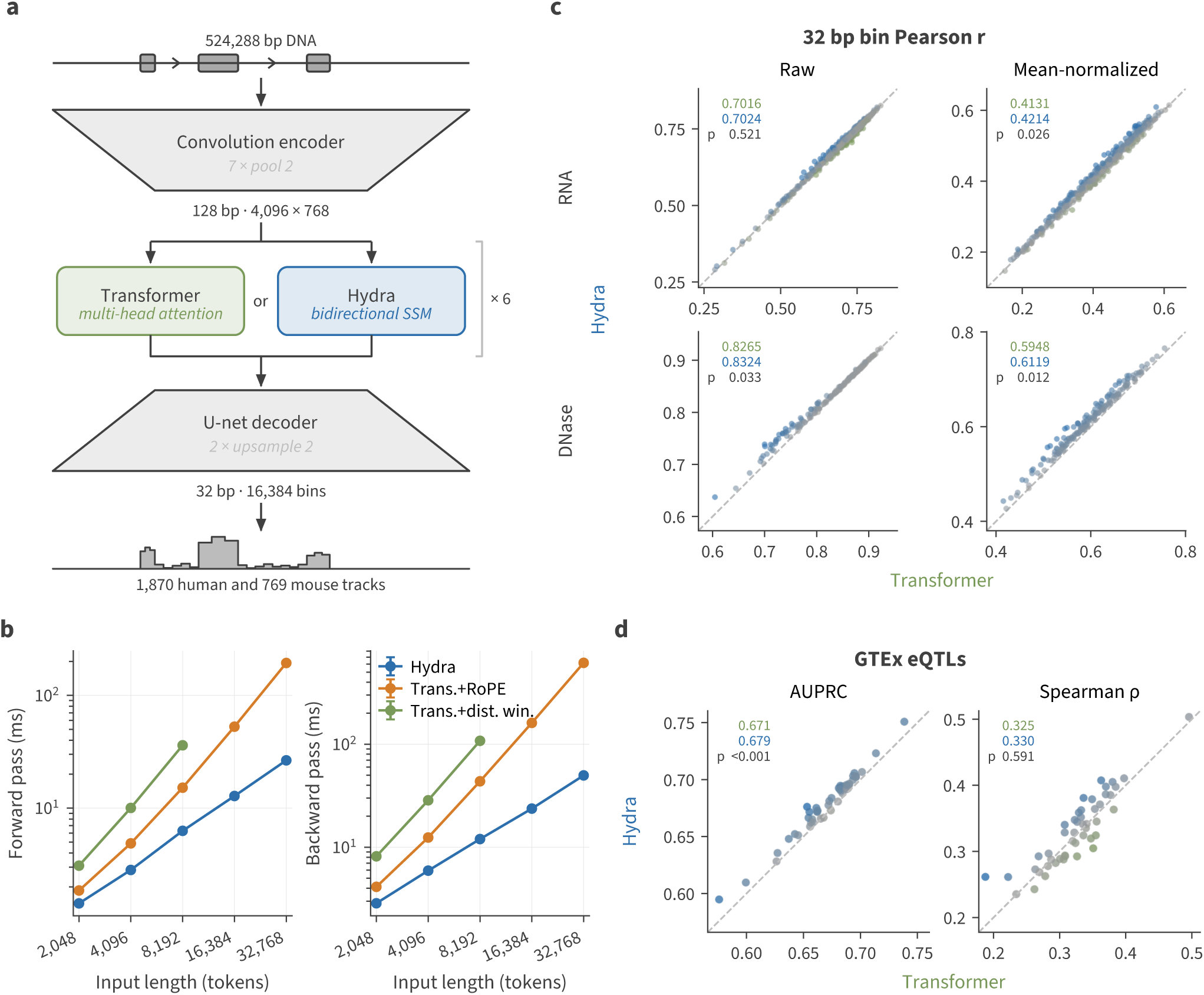
Hydra blocks are faster than attention and more accurate on held-out coverage and fine-mapped eQTLs. Every comparison holds the architecture fixed—convolution encoder to 128 bp resolution, six long-range blocks, U-net decoder to 32 bp coverage, and separate human and mouse heads predicting 1,870 and 769 tracks—and substitutes only the long-range block. The transformer biases attention with nested distance windows of geometrically increasing width, introduced with Enformer [16] and retained by Borzoi (Methods). **(a)** The shared architecture and the substitution the comparison isolates. **(b)** Forward (left) and backward (right) pass time of a single block against input length, at 1,024 channels, batch size 1, fp32, on one RTX 4090. Points are the mean of 20 timed iterations after 3 warmup iterations and error bars their standard deviation; backward time is the forward-plus-backward measurement minus the forward. The distance-window transformer block exhausts memory beyond length 8,192, so its line ends there. **(c)** Per-track Pearson *r* between predicted and measured 32 bp coverage on held-out test sequences, for RNA (top) and DNase (bottom) tracks, raw (left) and after quantile normalization and per-bin cross-track mean subtraction, which isolates specificity (right; Methods). Each point is one track, averaged over four replicates holding out different test folds, and is tinted toward the model that wins it. Dashed line, parity; the annotation gives each model’s mean over tracks and a two-sided paired *t*-test *p* over the four replicates (Methods). **(d)** The four-replicate ensemble of each family on GTEx v11 fine-mapped eQTLs (SuSiE PIP *≥* 0.9) against matched negatives, in 48 tissues. Both models score a variant with a single signed log fold change at its designated gene, as in Figs. 2 and 3; classification ranks by its absolute value. Left, AUPRC; right, Spearman *ρ* between signed predicted and measured effect size on positives. Each point is one tissue; parity line as in (c), and the annotation gives each model’s mean over tissues and a cluster bootstrap *p* over variants (Methods).

At 8,192 tokens, the longest input all three blocks fit in memory, Hydra blocks are 5.7-fold faster than transformer blocks that encode distance with log-spaced window features, and 2.4-fold faster than transformer blocks using rotary embeddings (RoPE) (Fig. 1b). The two transformer variants differ only in how they encode relative distance. The distance windows were introduced with Enformer [16] and retained by Borzoi (Methods). Consistent with others’ reports [17], RoPE transformers scored below distance-window transformers on 10 of 12 coverage, gene, and eQTL metrics, significantly only on raw DNase coverage (Supplementary Fig. S1). We therefore focus on distance-window transformer blocks.

Schedule-free Adam [18] scored above cosine decay on all four coverage metrics and four of six gene metrics, and tied it on eQTL classification (Supplementary Fig. S2). Schedule-free optimizers replace the learning rate schedule with a form of iterate averaging that interpolates between Polyak-Ruppert and primal averaging, evaluating gradients at a momentum-weighted combination of the base and averaged iterates. This removes the need to fix a training stopping time in advance.

Hydra predicts held-out coverage more accurately than the transformer on DNase tracks and on RNA tracks after the mean-normalizing transformation that isolates specificity (Fig. 1c). Gene-level accuracy favors Hydra in five of six comparisons, including cross-sample prediction over the most variable genes, though none individually reaches significance across four replicates (Supplementary Fig. S3). Hydra also classifies fine-mapped eQTLs against their matched negatives more accurately, 0.679 versus 0.671 AUPRC, higher in 46 of 48 tissues (p < 0.001; Fig. 1d). Effect-size correlations were 0.330 and 0.325 Spearman ρ, respectively (p = 0.59).

Hydra’s linear scaling let us drop convolution/U-net pairs one at a time, running the blocks on finer-resolution and therefore longer sequences. With fewer convolution/U-net pairs, parameter count decreased from 36.1 M (128 bp / 2 U-net) to 32.8 M (64 bp / 1 U-net) and 30.0 M (32 bp / 0 U-net). Both training and validation metrics rose as the U-net blocks came out, despite the smaller parameter count (Supplementary Fig. S4). As these models operate Hydra blocks on longer sequences, training time increases from 60.5 min/epoch (128 bp / 2 U-net) to 67.1 min/epoch (64 bp / 1 U-net) and 81.0 min/epoch (32 bp / 0 U-net).

Two further variations failed to improve these models. Hybrid architectures interleaving transformer and Hydra blocks, which help in language modeling [19], trained less stably here and scored lower. Varying the SSM state dimension, central to Mamba-2’s gains over Mamba-1 [7, 13], changed accuracy little, so we fix it to one for all reported models.

### 2.2 Cerberus foundation ensemble

These results led us to scale a convolution-SSM architecture with schedule-free optimization to full leading-model size. First, we refined the Borzoi Prime track collection [20], which added many scRNA-seq cell type pseudobulk profiles to the original Borzoi collection. We further added CLIP-seq profiles from ENCODE and Goodarzi lab [21], polyadenylation regulator perturbation scRNA-seq pseudobulk profiles [22], and 4sU metabolic labeling and actinomycin D transcription shutoff RNA-seq time courses that measure mRNA decay. Due to Hydra’s favorable memory scaling, we increased input sequence length to 786 kb.

We trained a 138.3 M-parameter model, against 185.9 M for Borzoi, with a tower of convolution and pooling blocks to 1536 channels where each position represents 32 bp from the original sequence. We replaced Borzoi’s transformer tower with a stack of 8 Hydra blocks. We trained 8 replicate models, each holding out a different test set. We refer to the ensemble of these eight models as Cerberus (Fig. 2a).

**Fig. 2.**
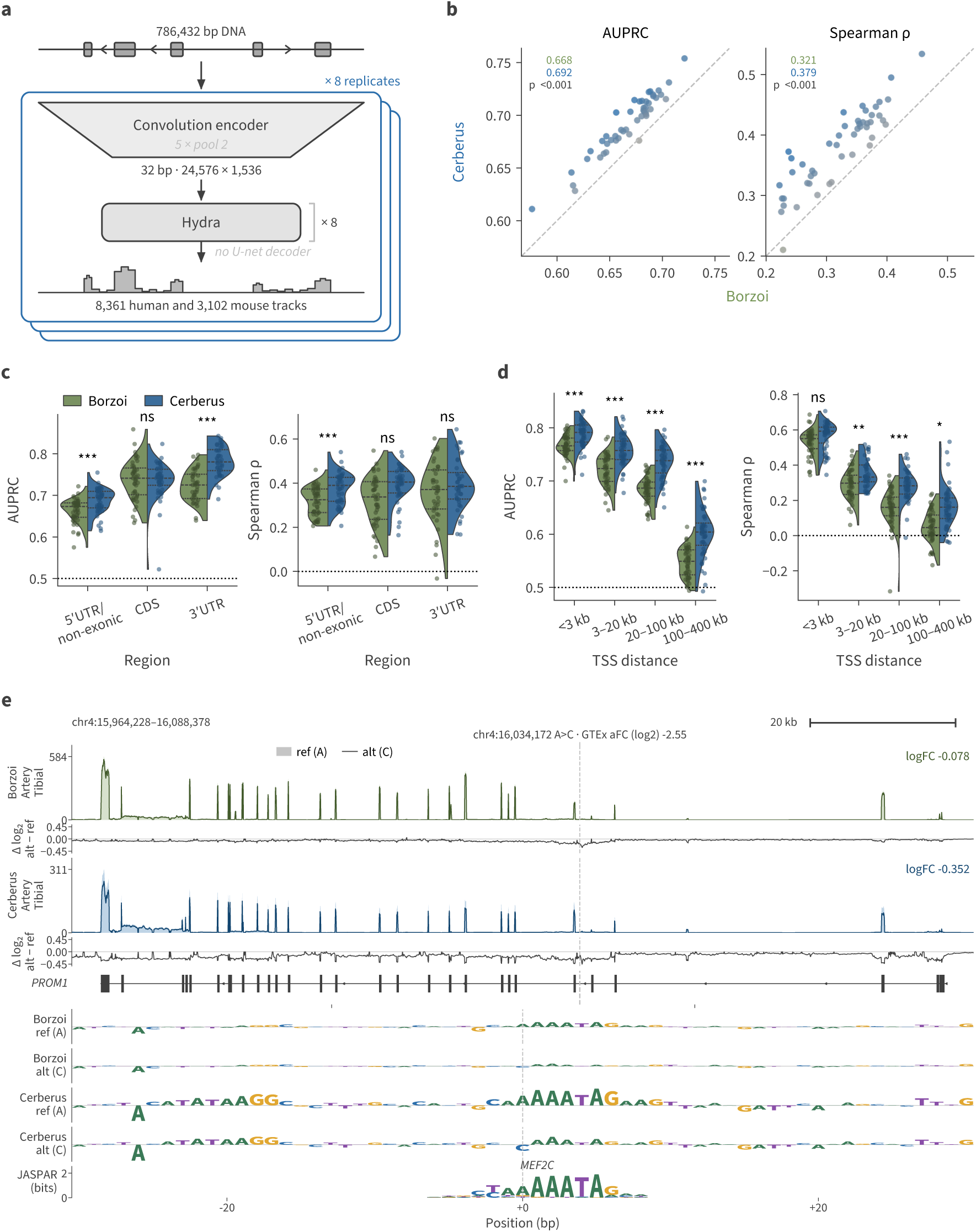
Cerberus classifies fine-mapped eQTLs and estimates their effect sizes more accurately than Borzoi. **(a)** The Cerberus architecture. A convolution encoder reduces 786,432 bp of input sequence to 32 bp bins, eight Hydra blocks mix at that resolution without a U-net decoder, and separate human and mouse heads predict 8,361 and 3,102 coverage tracks. Cerberus is the mean of eight such replicates, each holding out a different fold. **(b)** Per-tissue comparison of Borzoi and Cerberus on GTEx fine-mapped (PIP *≥* 0.9) eQTLs. Each point is one tissue; axes give Borzoi (x) versus Cerberus (y). Left, AUPRC for discriminating causal eQTLs from matched negatives; right, Spearman *ρ* between the signed predicted log fold-change and the measured eQTL effect size on positives. Points are tinted toward the winning model, the dashed line marks parity, and the annotation reports each model’s mean and the cluster bootstrap *p* over variants. **(c)** The same AUPRC (left) and Spearman *ρ* (right) metrics stratified by the eQTL’s genomic region (5*^′^* UTR/non-exonic, CDS, 3*^′^* UTR). **(d)** The same two metrics as a function of distance from the variant to the eQTL gene’s TSS, binned in base pairs. Cerberus’s advantage is largest for distal variants. **(e)** A fine-mapped tibial-artery eQTL for *PROM1* (chr4:16,034,172 A*>*C), intronic and 41.7 kb from the TSS. Top, predicted reference (fill) and alternate (line) RNA coverage under Borzoi and Cerberus, each above its per-bin log_2_(alt + 1) *−* log_2_(ref + 1); at right, the gene-level logFC. Bottom, in-silico mutagenesis (ISM) over *±*30 bp of the same statistic, both models on both allelic backgrounds and sharing a *y*-axis, with the matched MEF2C recognition matrix aligned beneath. Dashed line, variant position. In (c) and (d), split violins show per-tissue point estimates for Borzoi and Cerberus with every tissue overlaid as a point, and stars mark a cluster bootstrap over variants (*^∗^ p <* 0.05, *^∗∗^ p <* 0.01, *^∗∗∗^ p <* 0.001). See Methods for full details on the violin, inclusion, and testing conventions.

Differences in dataset preparation—fold division and coverage-track transformations—preclude a perfectly clean comparison to prior models like Borzoi. Even so, on similarly transformed bin-level DNase prediction Cerberus is more accurate, particularly after mean normalization isolates tissue/cell-type specificity (Supplementary Fig. S5a). The differing bin-level transformations can be undone to assemble comparable gene-level predictions by reversing each transformation, summing across exons, and log-transforming. Evaluated on the 848 genes shared between the Borzoi test set and one of our held-out folds, Cerberus is more accurate, with the gain again largest for tissue/cell-type specificity (Supplementary Fig. S5b).

### 2.3 Fine-mapped QTL benchmark

Most applications of these models—scoring variants, inferring regulatory motifs, designing sequences with a chosen activity—depend on their assigning predicted activity to the bases that cause it. Population genomic profiles measure that assignment directly, and since none of the models benchmarked here trains on genetic variation, the comparison puts them on the same footing. As above, we evaluated the models using a fresh GTEx v11 fine-mapped eQTL analysis. Linkage disequilibrium prevents most eQTLs from being resolved to a single causal variant, but SuSiE fine-mapping has now been applied with consortium rigor across the GTEx v11 eQTL, splicing QTL (sQTL), and polyadenylation QTL (paQTL) computations in 48 tissues, yielding positives we can treat as high-confidence.

The same fine-mapping supplies negatives, drawn from significantly associated variants that SuSiE assigns PIP below 0.01 in every tissue. Left uncontrolled, four properties separate fine-mapped positives from unselected variants without predicting their regulatory effects: transcript region overlap, distance to the transcription start site (TSS; or splice/polyadenylation site), minor allele frequency, and gene expression level. We therefore matched each positive to a negative on exact alleles, allele frequency, and gene expression, with region matching for eQTLs and distance matching for 5*^′^* UTR/non-exonic eQTLs, sQTLs, and paQTLs (Supplementary Fig. S6, S7, S8; Methods). Taking the union across tissues, 23,836 eQTL, 20,732 sQTL, and 4,991 paQTL SNPs (i.e. substitutions) comprise the positive sets, and each was matched to a negative SNP. A fine-mapped QTL is called separately in each tissue it acts in, so these sets hold 58,122 eQTL, 58,517 sQTL, and 14,823 paQTL variant–tissue pairs, but a variant’s measurements correlate strongly across the tissues it recurs in (intraclass correlation 0.83 for eQTLs), as do its predictions (0.88 to 0.96; Supplementary Fig. S9). Thus we compare models throughout with a cluster bootstrap over variants, each resampled with all of its tissues. Small indel collections are attainable via the same strategy, but we declined to do so here due to the complexity of indel scoring with these models [23]. Positive and negative QTLs in these sets come matched to a gene, and we recommend gene-specific scores for model evaluation (and caution against gene-agnostic scores because regional gene density was not controlled for).

### 2.4 Comparison with Borzoi

We scored every variant with the Borzoi and Cerberus ensembles using a log fold change statistic: for each variant–gene pair we sum the predicted coverage over output bins overlapping the gene’s exons and take the log ratio between the alternative and reference allele sequences. Across tissues, Cerberus separates fine-mapped eQTLs from their matched negatives more accurately than Borzoi, 0.692 versus 0.668 mean AUPRC, and estimates their allelic fold change better, 0.379 versus 0.321 mean Spearman ρ (Fig. 2b). It leads in 47 of 48 tissues on both metrics, and classification improves in every region and distance stratum except coding sequence (Fig. 2c,d). Transcript-overlapping variants in the CDS and 3*^′^* UTR are classified with greater accuracy than the remainder (0.741 and 0.780 versus 0.694 AUPRC), although their effect sizes are estimated no better. Concordance between model and QTL falls with TSS distance, from 0.792 AUPRC and 0.593 ρ within 3 kb to 0.604 and 0.161 beyond 100 kb. The Cerberus advantage moves the other way, widening from +0.022 to +0.056 AUPRC and from +0.044 to +0.099 ρ over the same span.

Fine-mapped positives differ in how strongly the association data support them. We used the GTEx metadata to decompose model accuracy along axes of statistical confidence (Supplementary Fig. S6). Both discrimination of causal from matched non-causal variants and agreement between predicted and measured allelic fold change increase with SuSiE PIP and eQTL significance. Thus, some disagreement may reflect errors in the QTL labels rather than in the model predictions.

We selected an example eQTL by ranking on Cerberus’s improvement over Borzoi in predicted effect size and on distance to the TSS. Cerberus predicts a 4.5-fold larger gene-level effect than Borzoi for a fine-mapped tibial-artery eQTL, intronic and 41.7 kb from the *PROM1* TSS (Fig. 2e). In the Cerberus ISM attribution, the variant falls in an A/T-rich motif matching the MEF2C recognition matrix, which the alternate allele disrupts; Borzoi attributes less to the same motif. The variant therefore likely acts by disrupting MEF2 binding. MEF2C is a plausible candidate in artery, where endothelial MEF2C inhibits smooth-muscle migration into the intima [24].

We continued these analyses to focus on classification of GTEx fine-mapped sQTL and paQTL causal variants from the negative sets that control for MAF, gene expression, and distance to splice junction or polyadenylation site. To measure predicted changes in within-gene coverage profiles, we scored each variant at its designated gene by the maximum profile shift (MPS). MPS is the largest absolute per-bin difference between reference and alternate log-coverage profiles after normalizing each to sum to one across the gene locus. The sQTL and paQTL evaluation sets display the same trend as eQTLs—accuracy increases with SuSiE PIP and QTL significance (Supplementary Fig. S7, S8).

Cerberus classifies both phenotypes more accurately than Borzoi, with the higher AUPRC in every tissue on sQTLs and 46 of 48 on paQTLs (means 0.718 vs. 0.689 and 0.709 vs. 0.691; both p *≤* 0.001; Supplementary Fig. S10a, c). The gain concentrates within 10 kb of the affected splice or poly-A site, where both models are most accurate and Cerberus leads in every distance bin by 0.014 to 0.036 AUPRC (all p < 0.05; Supplementary Fig. S10b, d). From 10 to 100 kb the two models differ by at most 0.010, and beyond 100 kb neither beats chance, so we excluded those positives, which concentrate in the least-powered tissues, from this comparison.

### 2.5 Comparison with AlphaGenome

AlphaGenome is a recently released sequence-to-function model that follows the general Borzoi framework of predicting aligned read coverage tracks directly from DNA sequence, while extending it along four axes: a longer 1 Mb input, base-pair output resolution, more parameters, and a wider set of modalities and objectives, including Hi-C contact maps, splice-junction classification, and junction-spanning read counts [2].

We compared Cerberus with AlphaGenome on fine-mapped GTEx v11 eQTLs, scoring both models with a single signed log fold change at the variant’s designated gene. We focused on expression because AlphaGenome’s API scores splicing and polyadenylation only with scorers defined against a gene’s annotated junctions or cleavage sites, in place of the annotation-free MPS we apply to Borzoi and Cerberus, and those scorers return no value for 22% and 48% of variants at their designated gene (Methods). Aggregated over all variants in a tissue, AlphaGenome is the more accurate model (Fig. 3a): it attains higher classification AUPRC in 40 of 48 tissues (means 0.703 vs. 0.692; p < 0.001).

**Fig. 3.**
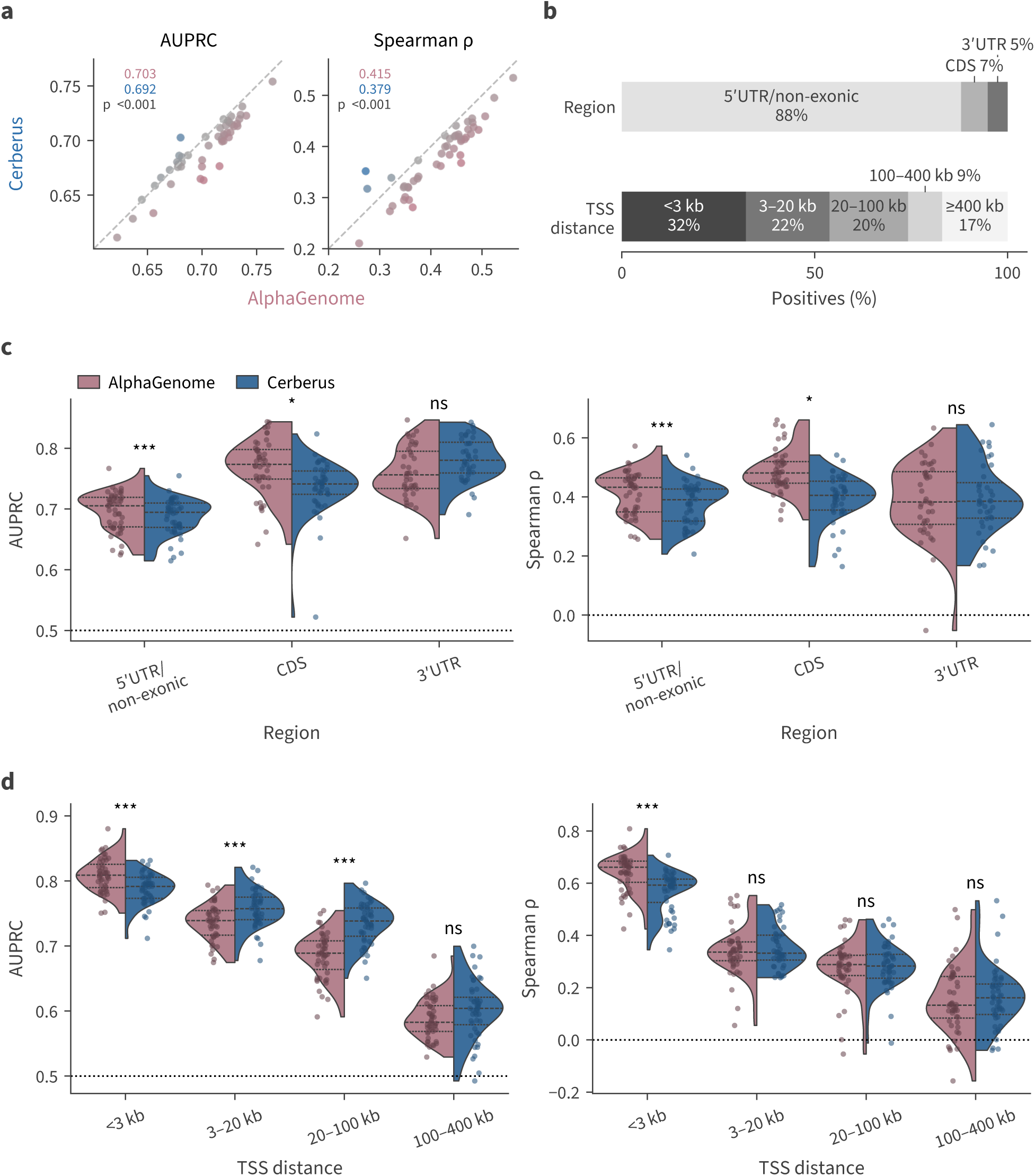
Cerberus and AlphaGenome have complementary strengths across eQTL classes. Models evaluated on fine-mapped GTEx v11 eQTLs (SuSiE PIP *≥* 0.9) in 48 tissues, each positive paired with a matched negative variant. Both models score a variant with a single signed log fold change at its designated gene; classification ranks by its absolute value, and effect-size correlation is Spearman *ρ* between signed predicted and measured effects on positives. **(a)** Per-tissue AUPRC (left) and Spearman *ρ* (right); each point is one tissue, and inset values are means across tissues with the cluster bootstrap *p* over variants. **(b)** Share of the variant–tissue positives held by each stratum of (c) and (d). Positives more than 400 kb from the TSS are counted here and dropped from (d). **(c)** The same two metrics stratified by the variant’s annotated region relative to its eQTL gene: coding sequence (CDS), 3*^′^* UTR, and the 5*^′^* UTR/non-exonic class, matched on distance to the TSS. **(d)** The same two metrics stratified by distance from the variant to the eQTL gene’s TSS. In (c) and (d), split violins show per-tissue point estimates with every tissue overlaid as a point, and stars mark a cluster bootstrap over variants (*^∗^ p <* 0.05, *^∗∗^ p <* 0.01, *^∗∗∗^ p <* 0.001, ns not significant); strata with fewer than 20 positives in a tissue are dropped. See Methods for full details on the violin, inclusion, and testing conventions.

However, that single ordering averages over classes of variant on which the two models behave differently (Fig. 3c). AlphaGenome’s advantage is strongest in coding sequence, where it leads by 0.021 AUPRC (median across the 45 tissues with sufficient CDS positives, p = 0.020). In addition, on the 5*^′^* UTR/non-exonic variants that dominate the benchmark, AlphaGenome holds a small lead (+0.009, p < 0.001). On 3*^′^* UTR variants the median difference reverses, +0.019 toward Cerberus, but the cluster bootstrap over the stratum’s 895 variants does not separate it from zero (p = 0.13).

TSS distance places the same crossover on a continuous axis (Fig. 3d). Within 3 kb of the TSS AlphaGenome leads (+0.019 AUPRC, p < 0.001); beyond 3 kb the advantage crosses over to Cerberus and peaks at 20–100 kb, +0.018 at 3–20 kb and +0.049 at 20–100 kb (both p < 0.001). The outermost bin holds a +0.016 median that is not significant over its 2,875 variants (p = 0.19).

Two features of the benchmark’s composition explain how these class-level results coexist with the tissue-level aggregate (Fig. 3b). Cerberus’s only region-level lead, in the 3*^′^* UTR, covers 5% of the positives, whereas the 5*^′^* UTR/non-exonic class AlphaGenome leads covers 88%. Cerberus’s distance-level leads are larger, 42% of positives at 3–100 kb against 32% within 3 kb, but panel a pools every class into a single ranking, so a model’s scores must be comparable between classes as well as correctly ordered within each.

AlphaGenome’s coding-sequence lead concentrates in the variants fine-mapped across many tissues (Supplementary Fig. S11), which coding eQTLs are more often than other classes, 11.3% in 11 or more tissues against 3.6% of 5*^′^* UTR/non-exonic variants. AlphaGenome’s Spearman ρ lead is 0.025 over all 840 coding variants (p = 0.32) but 0.192 (0.072 to 0.324) over the 95 fine-mapped in 11 or more tissues. No finer coding annotation, whether consequence class, exon, or position within the coding sequence, separates the models consistently (Supplementary Fig. S11).

### 2.6 Inference cost

We measured forward-pass latency and peak GPU memory for Cerberus on one A100 80 GB and take AlphaGenome’s from a published benchmark on the same GPU under the same protocol [25], its non-commercial license not permitting us to run it outside of the API (Supplementary Fig. S12). At 1 Mb, AlphaGenome’s training length and the longest the published benchmark reports, a Cerberus forward pass takes 135.7 ms and 4.1 GB against 681.1 ms and 36.1 GB, 5.0-fold faster on 8.8-fold less memory. At its own training length of 786 kb Cerberus takes 101.5 ms and 3.2 GB.

Borzoi is the cheaper model per forward pass at the lengths these two are trained on. Cerberus runs its long-range blocks at 32 bp resolution and Borzoi at 128 bp, so on a matched 524 kb input the Hydra stack processes 16,384 tokens against the transformer’s 4,096. Cerberus costs 1.5-fold more at that length, 68.9 ms against 45.2 ms. Per token the Hydra block is the faster of the two (Fig. 1b), and the four-fold token count absorbs that advantage. The prediction heads are a second source of that cost. Cerberus emits 8,361 human tracks on all 16,384 output bins, against Borzoi’s 7,611 on the 6,144 bins that survive its crop. Both heads run inside the timed forward pass, so their 137 million and 47 million predicted values are part of the latencies above. Latency grows as L^0.98^ in sequence length L for Cerberus and L^1.40^ for Borzoi, so the two cross between 1.05 and 1.31 Mb; at 2.10 Mb Cerberus takes 267 ms and 7.6 GB against 403 ms and 14.3 GB for Borzoi.

Every number above is one forward pass of one network, and the three models are released with different numbers of networks. Borzoi comes as four replicates and Cerberus as eight, each trained on a different fold split, so scoring one allele runs four and eight networks; the eQTL comparisons above run all eight. AlphaGenome instead distills 64 all-fold teacher models into one released student model, which its authors report as competitive with a mean ensemble of separately pretrained models [2], so its cost is one network throughout. At that level the per-pass comparison reverses on latency but not on memory: at 1 Mb the eight Cerberus networks take 1.09 s against AlphaGenome’s 0.68 s and, run one at a time, hold a 4.1 GB peak against 36.1 GB. Distillation is a property of the training recipe rather than of the block, and we have not applied it to Cerberus; if it transfers, it would cut the eight networks to one.

### 2.7 Hydra block interpretation

Attention maps made the Borzoi transformer’s sequence mixer legible, its heads highlighting transcription start sites and other exon boundaries [1]. Cerberus replaces attention with Hydra blocks, whose sequence mixer, conditional on the input-dependent parameters, is multiplication by a quasiseparable matrix [13, 14], so the same readout is available to 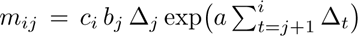 for *i* > *j*, with the reverse pass supplying *i* < *j*, so each head’s operator, the analogue of an attention map, follows from the parameters and the input without probing. The step size Δ*_t_* scales how much position t writes into the state. We summarize a head’s reach by the half-life of its decay at its median step size, the distance over which its contribution falls by half. We measured both in all eight Cerberus replicates, 1,280 heads over 8 blocks, on 16 held-out sequences of 786 kb (Methods).

Half-life rises with block depth to a maximum in the seventh of eight blocks (Fig. 4b), from a median of 0.9 kb in the first block to 79 kb in the seventh, an 87-fold span. The seventh block exceeds the first in 8 of 8 replicates. Over the same blocks the lower quartile of half-life moves from 0.3 to 1.9 kb and the upper quartile from 9.2 to 329 kb, so deep blocks retain their local heads and add long-range ones. The rendered operators show that trend directly (Fig. 4a), 94% of the median first-block head’s weight lying within 3 kb of the diagonal against 26% in the seventh. Within the seventh block, heads run from a pure local average to one placing 99% of its weight in a few vertical stripes (Supplementary Fig. S13a).

**Fig. 4.**
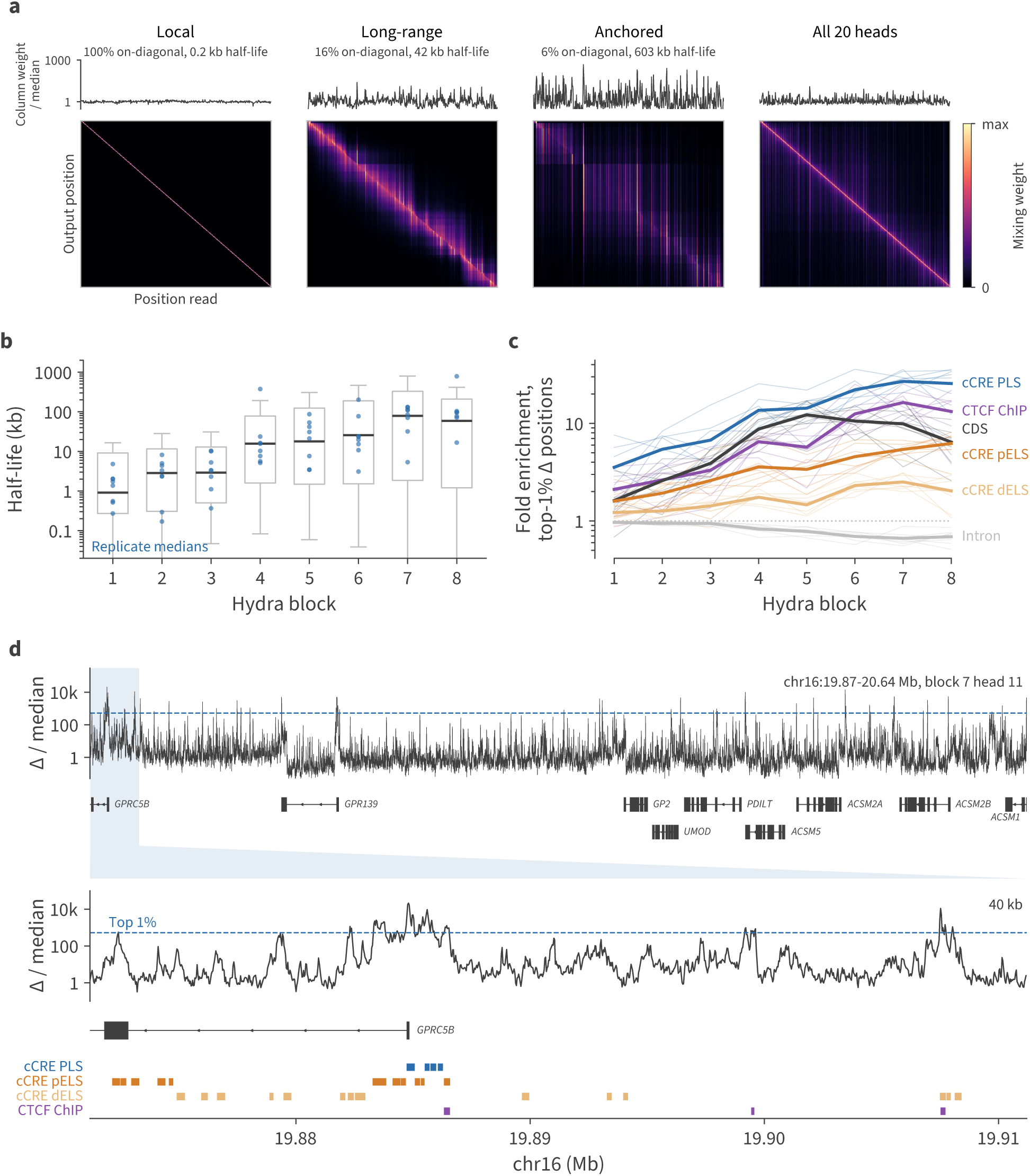
Hydra reach grows with depth, and the step size of deep heads spikes at regulatory elements. All panels cover the eight Cerberus replicates, 8 blocks *×* 20 heads each, evaluated on 16 held-out sequences of 786 kb. **(a)** Operators of three heads of the seventh block in replicate 1, descending in on-diagonal weight — the most diagonal head, a long-range head, and the head panel d tracks — and the sum over the block’s 20 heads. On-diagonal weight is the share of *|m_ij_ |* within 3,072 bp of the diagonal and is printed under each head’s name with its half-life. *|m_ij_ |* is taken on the held-out window of panel d. Tiles are drawn after trimming the recurrence warm-up at each edge, and each is scaled to its own 99.9th percentile, since the subsequent output projection absorbs the absolute scale of a head’s operator. Above each tile is its column weight, *|m_ij_ |* over the median of that quantity, the total a source position writes into the window. **(b)** Half-life of all 160 heads per block, boxes spanning the interquartile range with whiskers at 1.5 IQR, clipped to the axis, and outliers omitted; the eight replicate medians are overlaid. **(c)** Fold enrichment of each replicate’s anchors, its top 1% of positions by step size, on six annotation classes, relative to the class’s density in the scored windows; heavy lines are the mean over replicates and thin lines the individual replicates. **(d)** The step size of example head 11 of the seventh block in replicate 1, over a held-out window, as a multiple of its median. Top, the whole 770 kb window (chr16:19,870,822–20,640,870) on a log scale, with the head’s top-1% threshold at 523-fold dashed, over the protein-coding gene models of the window. The shaded span expands onto the tiers below, a 40 kb zoom on the strongest anchor, its gene model, and four annotation rows. The tallest anchor of the zoom reaches 21,000-fold the median on the promoter-like cCRE of *GPRC5B*; the second, 11,000-fold, sits 23 kb away on a CTCF ChIP-seq peak.

Those stripes are the positions at which the step size spikes. In the seventh-block head shown, Δ sits near its median for tens of kilobases along the window and exceeds it 1,000-fold at isolated positions (Fig. 4d). The strongest position falls in the promoter-like cCRE of *GPRC5B*, and the strongest outside that element on a distal CTCF ChIP-seq peak 23 kb away. The median first-block head puts 12% of its summed Δ in its top 1% of positions, and the median seventh-block head 40% (Supplementary Fig. S13b). Uniform Δ would put 1% there. The input gate b is shared across heads and is less concentrated across positions than Δ. With ngroups = d_state_ = 1, it is a single channel shared by all 20 heads of a block. At every depth it puts 1.1–1.4% of its total in its top 1% of positions. These concentration statistics identify Δ as the more spatially concentrated input factor.

Defining a head’s anchors as its top 1% of positions by Δ, we counted their overlap with ENCODE cCREs, CTCF ChIP-seq peaks, GENCODE transcription start sites, coding sequence and introns (Fig. 4c). The first block’s anchors are enriched 3.6-fold on promoter-like cCREs and 2.1-fold on CTCF ChIP-seq peaks, over each class’s background density in the scored windows. By the seventh block the same enrichments reach 27-fold on promoter-like cCREs and 16-fold on CTCF peaks, in 8 of 8 replicates (per-class p *≤* 5 *×* 10*^−^*^5^ against a circular-shift null; replicate ranges 12–36 and 10–25-fold). The 92% of CTCF peaks with no promoter-like cCRE under them enrich 15-fold, against 38-fold for those that do (Supplementary Fig. S13d), so the anchors mark CTCF sites in their own right. Anchor density peaks in the 32 bp bin holding the element midpoint, 29-fold for both classes, and falls to 9.9 and 2.2-fold 500 bp away (Supplementary Fig. S13e), so the anchors track the element rather than the region around it. GENCODE TSSs, 65% of which fall in a promoter-like cCRE, reach 37-fold by the seventh block (replicate range 16–49-fold), so an annotation built from gene models agrees with one built from chromatin. Enrichment declines with distance from the TSS, the three cCRE classes falling from 27-fold on promoter-like through 5.4-fold on proximal enhancer-like to 2.5-fold on distal enhancer-like. Coding sequence reaches its max 12.3-fold in the fifth block, peaking two blocks shallower than the regulatory classes, though short 5*^′^*UTRs leave part of that signal promoter-adjacent. The ordering sharpens at stricter thresholds — promoter-like cCREs 27-fold at the top 1%, 56-fold at 0.5%, 105-fold at 0.1% — and heads within the seventh block anchor at overlapping positions (median pairwise Jaccard 0.11, against 0.004 with one head of each pair shifted by a random offset), which is why significance is assessed per replicate rather than per head.

Many mammalian promoters lie in CpG islands, but base composition does not account for the promoter enrichment (Supplementary Fig. S13d). Splitting promoter and non-promoter bins at the median GC content of promoter-like cCREs leaves low-GC promoters at 18-fold, against 35-fold for high-GC promoters and 2.0-fold for high-GC non-promoter bins. Computing each anchor’s expected overlap from bins in its own GC decile still leaves promoter-like cCREs at 12-fold and TSSs at 17-fold enrichment.

Although the blocks are bidirectional, asymmetries emerge. Forward and reverse half-lives differ 6.3-fold within the median head, and 11% of heads are effectively one-directional, one pass’s output averaging under 1% of the other’s in magnitude (Supplementary Fig. S13c). That fraction is 47% in the first block and at most 3% in blocks 3 through 6. Of the 160 first-block heads across the eight replicates, 37 are forward-only and 38 reverse-only, so neither direction dominates.

## 3 Discussion

Sequence-to-function models predict a variant’s effect on a gene from DNA sequence alone, and genetic analyses now use them to prioritize candidate causal variants and propose mechanisms for them. Each generation of models has improved on the last, but substantial headroom remains. On fine-mapped GTEx eQTLs paired with negatives matched for basic variant properties, the best model in this comparison reaches 0.70 AUPRC against a chance rate of 0.5, and Cerberus classification falls from 0.79 AUPRC within 3 kb of the TSS to 0.60 beyond 100 kb. The data needed to close that gap, unlike language modeling’s corpus, cannot readily grow. New experiments add labels to the same reference chromosome sequences. They deepen the annotation of this fixed sequence collection without expanding the sequence diversity available for learning regulatory rules. Architecture, objective, and optimization offer ways to improve accuracy with the available data.

The architecture reported here is one such gain. Replacing Borzoi’s transformer blocks with bidirectional state space blocks improved accuracy and lowered the cost of the block together. At matched scale and on matched data, Hydra was the more accurate block across four training replicates, while running 5.7-fold faster at 8,192 tokens. The efficiency then bought a simpler architecture, because Hydra’s cost grows linearly with sequence length, so we could operate it on finer-resolution sequence and remove the U-net decoder entirely, arriving at a smaller convolution-SSM model with better validation accuracy than the one it replaced. Scaled to a full foundation ensemble, Cerberus improves on Borzoi across all three fine-mapped GTEx phenotypes. Against AlphaGenome, it trades leads by class of variant, ahead on variants 3 to 100 kb from the TSS and behind in coding sequence and within 3 kb. Regulatory annotation thins with distance from the gene, so distal variants are where model predictions add the most to variant prioritization.

State space models encode positional relationships in their recurrent dynamics rather than in explicit positional encodings. Transformers cannot. In the Enformer formulation and its successors, relative positional encodings must be added to inform the attention mechanism that nearby positions should interact more strongly than distant ones. Off-the-shelf encodings such as RoPE have trailed those bespoke distance windows in regulatory genomics models, in these experiments, in many previous unpublished attempts of ours, and in others’ work [17]. RoPE can be absorbed into the query-key product and the distance windows cannot, so fused attention kernels serve only the encoding that costs accuracy. SSMs build this inductive bias into their structure, because the recurrence naturally attenuates influence with distance, at a rate that varies across heads and adapts to the input through the selective parameterization. This matches the distance-dependent logic of gene regulation, where the influence of a regulatory element on a promoter generally decays with genomic distance but in a context-dependent manner shaped by intervening elements such as insulators. Attention maps from the original Borzoi transformer also highlight exon boundary elements—transcription start sites, splice junctions, and polyadenylation sites—as prominent features, and these gene structure landmarks are likewise well suited to the SSM’s selective gating mechanism. Work comparing SSMs and transformers in the language domain has found that SSMs tend to be more data-efficient, with the primary exception being tasks that require precise associative recall over long contexts [26]. Such recall requires the state to hold arbitrary content, because the query that will select it arrives only at the end. In contrast, regulatory elements and gene structure landmarks are apparent as the model encounters them, and the state holds only a few types.

What the trained Hydra blocks compute resembles the attention maps of the transformer blocks they replace. Reach is graded by block depth, and the deepest heads write into the SSM memory most strongly at transcription start sites, promoters and CTCF peaks, and more weakly at enhancers with distance from the TSS. Our interpretation analysis identified two forms of specialization in the first block that motivate architectural ablations. Its heads mix over short distances, and nearly half of them read in only one direction. A narrower first block or a penalty on directional asymmetry could test whether these specializations improve prediction or leave capacity unused.

We emphasize molecular QTL evaluations deliberately. Comparing sequence-to-function models on held-out sequence requires matching their training tracks, preprocessing, and train/test splits (Supplementary Fig. S5), which binds each new model to its predecessor’s data choices when those choices merit exploration of their own. It is also the wrong target, because held-out accuracy is best understood as a guard against memorization rather than as an application, and the uses that motivate these models do not ask for predictions in regions of the genome for which no data exist. Those uses ask instead for causal attribution within the genome: how the sequence surrounding a gene determines its expression, how mutation alters that expression over evolution, and how variation across a population alters expression. The last of these connects these models to trait biology and to the interpretation of genetic associations. Population genomic profiles measure that relationship directly, on an axis external to any model’s training choices, and that independence recommends them as the shared basis for comparison. We therefore refreshed our prior GTEx evaluation rather than adopting an existing one, matching each positive variant to a negative on four properties that either raise the power to detect an association or are readable from annotation alone. Matching controls for these properties, sharpening the comparison of sequence-based regulatory predictions. Fine-mapped QTLs are themselves an inference, not a ground truth. A QTL call is the output of a long analytical pipeline with many defensible but sensitive settings. Variants fine-mapped in one consortium frequently fail to replicate in another, and the shortfall is not confined to marginal calls. A third of GTEx blood credible sets resolved to a single variant at PIP > 0.95 have nothing recovered for the same gene in TOPMed, a rate that tightening the threshold to 0.99 barely improves and that reaches 47% for lymphoblastoid lines assessed against MAGE [27]. Replication failure overstates error, since the replication set has false negatives of its own, but the same study puts the non-causal fraction among these high-confidence variants at 21–24%—still far above what a PIP of 0.95 asserts. Our own results carry that signature. In all three phenotypes, agreement between model and QTL rises with both SuSiE PIP and association significance, as expected if the weaker-evidence positives include variants that are not causal. We therefore read disagreement symmetrically, because where model and QTL diverge, the error is not always the model’s.

Model evaluation against QTL sets is more productive as a loop than as a leaderboard. Carefully constructed QTL sets identify better models and steer their development. In turn, models with demonstrated accuracy in a given regime can inform the analytical choices—variance models, inferred factor correction, fine-mapping priors—that QTL analyses presently make on discovery count alone. That second direction is already underway [3, 28]. Model scores predict which high-confidence variants fail to replicate better than PIP does, evidence that the models carry information the association statistics lack [27]. Folding those scores into the fine-mapping posterior ranks the replicating variant first in its credible set 79% of the time, against 67% without them [27]. Such use demands care, because the models are not uniformly reliable across regimes. Accuracy falls with distance to the affected site in all three phenotypes, so a prior built from model scores would be tighter on proximal variants than on distal ones. Single-cell QTL mapping multiplies these analytical choices, starting with whether to pseudobulk cells or model them directly.

We release the Cerberus models and training code as an open-source PyTorch implementation [29], departing from the TensorFlow codebase in which we developed Basenji and Borzoi. Research into efficient sequence-mixing blocks now appears predominantly in PyTorch, so a new block can be tested without a TensorFlow reimplementation. Borzoi has already been extended in PyTorch for accelerated inference [17], single-cell multiomic profiles [30], and cell type- and disease-specific expression [31]. Tools for sequence design [32] and for attribution and motif discovery [33] belong to the same software ecosystem, so Cerberus and future models built on its code compose with them directly.

## 4 Conclusions

For researchers training long-range sequence-to-function models, the bidirectional state space mixer is a practical alternative to self-attention. At matched scale and on matched data, models built on it were more accurate than transformers using either log-spaced distance windows or RoPE, and the block itself ran faster than both. Linear scaling with sequence length let us run the block at 4-fold finer resolution, which removed the need for a U-net decoder. We trained Cerberus, a foundation ensemble that predicts coverage and attributions across 8,361 human and 3,102 mouse functional genomics tracks, including cell-type-resolved pseudobulks from several single-cell atlases. Fine-tuning adapts these weights to new data [15].

For researchers studying genotype–phenotype relationships, Cerberus scores variant effects on expression, splicing, and polyadenylation more accurately than Borzoi, and trades leads with AlphaGenome by class of variant. Those comparisons rest on a GTEx v11 benchmark whose matched negatives help separate sequence-based inference from signal basic annotations supply, released with the code that constructs it [34]. Distal regulation remains the hard case for every model in this comparison, and these sets will help measure whether the next generation closes that gap.

## 5 Methods

### 5.1 Reduced search dataset

To explore model training hyperparameters, we constructed a dataset that enabled reasonably sized models to train in several days on an A100 GPU. At a high level, we followed Borzoi and Borzoi Prime dataset construction [1, 20]. We extracted 524 kb DNA sequences strided by one quarter of that length, and aggregated functional genomics data into 32 bp bins. We transformed these tracks as in prior work: ENCODE DNase and ChIP tracks are log fold changes, and the remaining tracks are summed counts, which we square-root to compress extreme values.

We downsampled these tracks randomly by ^1^ to a total of 1870 human and 769 mouse tracks. The human tracks consist of 199 ENCODE DNase, 275 ENCODE histone modification ChIP-seq, 459 ENCODE RNA-seq, 89 GTEx RNA-seq, 368 FANTOM CAGE, 264 Tabula Sapiens 3*^′^* scRNA-seq cell type pseudobulks, 95 Tabula Sapiens SMART-seq cell type pseudobulks, and 121 CATlas snATAC-seq cell type pseudobulks. The mouse tracks consist of 20 ENCODE DNase, 120 ENCODE histone modification ChIP-seq, 9 ENCODE ATAC-seq, 120 ENCODE RNA-seq, 194 FANTOM CAGE, 54 Tabula Muris 3*^′^* scRNA-seq cell type pseudobulks, 112 Tabula Muris SMART-seq cell type pseudobulks, 117 CATlas snATAC-seq cell type pseudobulks, and 23 sci-ATAC-seq atlas cell type pseudobulks. These sources are a subset of those cited below for the full foundation dataset.

We linked homologous human and mouse sequences via whole genome alignments and divided the sequences into ten folds [35]. For each replicate, we trained on eight folds and used the remaining two for validation and test. We trained models for a maximum of 80 epochs and chose the final parameters based on the epoch with the best validation Pearson r averaged across all tracks.

### 5.2 Full foundation dataset

To form a more comprehensive dataset for a foundation model, we extended the sequence length to 786,432, also striding by one quarter that length and dividing into 16 folds with homologous human and mouse sequences linked as above [35].

The human tracks consist of 2,028 RNA-seq, 1,741 3*^′^* RNA-seq, 1,678 CAGE, 1,282 ChIP-seq, 674 DNase-seq, 496 CLIP-seq, and 462 ATAC-seq. The mouse tracks consist of 953 RNA-seq, 720 CAGE, 562 ATAC-seq, 560 ChIP-seq, 214 3*^′^* RNA-seq, and 93 DNase-seq. Bulk RNA-seq, DNase-seq and ChIP-seq come from ENCODE [36] and Roadmap Epigenomics [37], with further RNA-seq studies drawn through recount3 [38], among them GTEx [39]. CAGE comes from FANTOM5 [40], and the 402 human single-cell CAGE pseudobulks from an atlas of transcribed cis-regulatory elements [41]. Transcriptomic cell type pseudobulks come from Tabula Sapiens [42], Tabula Muris [43], and an adult human brain atlas [44]. Accessibility cell type pseudobulks come from CATlas [45–49] and a mouse sci-ATAC-seq atlas [50]. CLIP-seq comes from ENCODE and the Goodarzi lab [21], perturbation pseudobulks from a polyadenylation regulator screen [22], and pooled poly(A) site profiles from PolyA DB [51]. mRNA stability time courses, by 4sU metabolic labeling or actinomycin D transcription shutoff, come from studies drawn through recount3 [38]. Every track is listed in Supplementary Table S1with its assay, source, sample, and accession.

### 5.3 State space model sequence mixers

While self-attention enables direct pairwise interactions between all positions, it incurs quadratic computational cost in the sequence length. We investigated replacing the transformer blocks with blocks based on structured state space models (SSMs), which achieve linear-time sequence mixing.

SSMs draw on the classical framework of linear dynamical systems, discretized to operate on sequences. Given an input sequence **x** = (*x*_1_, . . ., *x_L_*), an SSM computes an output through the recurrence *h_t_* = *A_t_h_t−_*_1_ + *B_t_x_t_*, 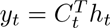, where *h_t_ ∈ R^N^* is a latent state and *A_t_, B_t_, C_t_* are learned parameters that may depend on the input at position t. This input-dependent (“selective”) parameterization, introduced in Mamba [7], allows the model to modulate how it accumulates and retrieves information as a function of the sequence content, while retaining a recurrent structure computable in *O*(*L*) time. Mamba-2 [13] restricts *A_t_* to a scalar times the identity, under which the recurrence can be expressed as multiplication by a semiseparable matrix *M*, where each entry 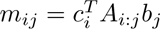 for *i* ≥ *j* and *m_ij_* = 0 otherwise, and computed by a chunked algorithm, structured state space duality (SSD), built on matrix multiplications. However, this lower-triangular structure enforces causality: each output depends only on current and preceding inputs. This unidirectional constraint is unnecessarily restrictive for gene expression prediction, where regulatory elements may lie upstream or downstream of their targets.

To construct a bidirectional SSM, a common heuristic is to run two independent SSMs—one forward, one backward—and combine their outputs by addition or concatenation [12, 52, 53]. Hydra [14] provides a more principled alternative grounded in quasiseparable matrices, which extend semiseparable matrices to include an upper-triangular component and a free diagonal:

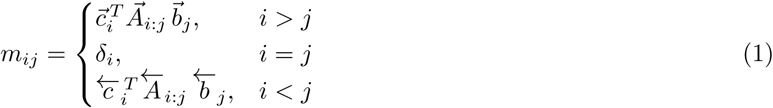

The lower- and upper-triangular blocks capture forward and backward sequence dependencies, respectively, while the diagonal terms δ*_i_* are parameterized as a learned, data-dependent function of the input, independent of the off-diagonal SSM parameters. This independence is the source of Hydra’s increased expressivity relative to naive bidirectional SSMs, where the diagonal is implicitly constrained by the vectors used to construct the off-diagonal entries. In practice, the quasiseparable matrix–vector product decomposes into two semiseparable operations on the forward and time-reversed input plus the diagonal term, allowing Hydra to reuse the efficient chunked-scan algorithm of SSD with minimal overhead. Forward and backward parameters share a single input projection, so the parameter count increases only marginally relative to a unidirectional model.

We constructed residual Hydra blocks consisting of RMS normalization followed by the Hydra sequence mixer, and substituted these for the transformer blocks in Borzoi’s tower.

### 5.4 Model and optimization parameters

Search models applied a convolution block with kernel width 13 and channels 384, followed by 6 convolution blocks that increase channel number to 768. Convolution blocks apply batch normalization, silu activation, and a width 5 convolution before max pooling. After the convolution tower, each position in the sequence represents 128 bp.

Subsequently, we applied a tower of 6 either transformer or Hydra blocks. Transformer blocks use 8 head self-attention, 2*×* expansion for the MLP, and dropout probability 0.05 before residual add. Attention is biased by 32 relative position features: 16 nested indicator windows centered on the query, of width (*L* + 1)^*k*/16^ − 1 positions for *k* = 1 . . . 16, and 16 sign-weighted copies that distinguish upstream from downstream. At the block’s *L* = 4,096 positions of 128 bp the widths grow by a factor of 1.68 each, so resolution is uniform in log distance. A learned projection maps the 32 features to per-head key vectors, added to the attention logits alongside learned content and position biases. Enformer introduced this family under the name `central_mask` and concatenated it with two others; its ablations found it sufficient, Borzoi retained it alone, and that is the formulation used here. Hydra blocks use 12 heads, 2*×* expansion, state depth 1, and dropout probability 0.05 before residual add.

Finally, we applied two U-net decoder blocks that each upsample the low-resolution trunk representation twofold and fuse it with the matching higher-resolution encoder skip. See below for additional description of U-net modifications with respect to Borzoi.

We trained using AdamW, batch size 4, 0.01 weight decay (100*×* reduced for normalization and final prediction layers), and global norm gradient clipping at 1.0. We tuned learning rate, momentum, and weight decay for each block type by manual iterative search, under both schedule-free [18] and cosine decay schedules. During training, we perform data augmentation by choosing a sequence shift from -1, 0, or 1 bp uniformly and reverse complementing the sequence (and reversing the tracks) half the time.

#### 5.4.1 U-net decoder block

We modified Borzoi’s U-net decoder block after comparing it with AlphaGenome’s. The Borzoi block normalizes and activates each input before merging. The coarse map x then passes through a pointwise convolution and is nearest-neighbor upsampled, while the skip map u is projected to the working width by a pointwise convolution. The two are summed and refined by a depthwise convolution, a pointwise convolution, and dropout, with no normalization before the refinement and no skip around it. Our new block makes two changes. First, the coarse pathway drops its normalization, activation, and convolution, so x is only rescaled by a learnable scalar (initialized to 0.9) and upsampled, letting the block learn how strongly to inject the trunk signal. Second, a normalization–activation is applied to the merged sum before the depthwise-separable refinement, and the refinement is wrapped in a residual connection, so the block adds a learned correction to the merged features instead of replacing them. The skip pathway is unchanged. Together these keep the block near an identity-plus-skip mapping at initialization. See Supplementary Fig. S14 for block schematic visualization.

#### 5.4.2 Spatial position-weighted loss

As in Borzoi, we decompose the coverage loss into a Poisson term on the total count over the region (magnitude) and a multinomial cross-entropy term on the normalized profile (shape), balanced by a weight λ [1]. Unlike Borzoi, which scored only a cropped central window, we compute the loss over the entire predicted sequence, so every output position is supervised. Borzoi’s cropped output bins received no direct supervision, which in our experience left predictions there unreliable.

Because boundary bins have a truncated receptive field and lack context beyond the input window, we down-weight them with a smooth super-Gaussian (flat-top) window rather than cropping. For an output bin at signed distance *u_i_* from the center (with *u_max_* = (*L* − 1)/2),

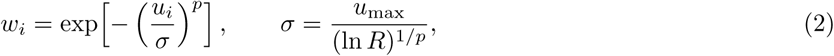

which holds *w_i_ ≈* 1 across a flat interior plateau and rolls off to exactly 1/R at the extreme edges. The exponent p sets the plateau flatness and *R* the center-to-edge dynamic range; we use *p* = 8 and *R* = 10 (edges weighted 0.1*×* the center). The weights multiply both loss terms per position, and both terms are divided by Σ*_i_ w_i_* so the objective stays a weighted mean independent of *p*, *R*, or sequence length. See Supplementary Fig. S15 for weight profile visualization.

#### 5.4.3 Cerberus

The full final model ensemble, which we refer to as Cerberus, begins with a 768 channel, 13 kernel width convolution block, followed by 4 convolution blocks that increase channel number to 1536. After the convolution tower, each position in the sequence represents 32 bp. Subsequently, we applied a tower of 8 Hydra blocks, using 5/3 expansion factor to 2560 channels, 20 heads, state depth 1, and dropout probability 0.08 before residual add.

We trained using AdamW, batch size 2, 0.003 learning rate, 0.015 weight decay (10*^−^*^4^ for normalization and final prediction layers), and global norm gradient clipping at 1.0. We tuned these hyperparameters roughly with several smaller replicate training runs, selecting on validation r. We trained 8 replicates for 50 epochs, each on 14 of the 16 folds with one held out for validation and one for test, so that 8 of the 16 folds serve as a test set. Each replicate trained for 11 days on one A100 GPU and kept the parameters from the epoch with the best validation Pearson r averaged across tracks.

Exact model specification and training recipes are available via the Baskerville repository [29].

### 5.5 Evaluation metrics

#### Coverage accuracy

For each held-out sequence, the model predicts binned coverage for every track. We evaluate each track *t* by pooling predictions and measurements across all sequences and bins and computing the Pearson correlation *r_t_* and explained-variance 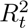.

#### Cross-track specificity

Track-level correlation is dominated by the shared coverage profile common to all assays. To isolate the *relative* differences between same-assay tracks, we evaluate specificity within each target group. Stranded pairs are first summed to a single total-signal track. Within a group, each track’s coverage vector over all positions is quantile-normalized to a common reference (the per-rank mean of the sorted tracks in the group). At each position the cross-track mean is subtracted, and per-track Pearson r is computed between the normalized predictions and measurements.

#### Gene expression accuracy

We form strand-specific, non-overlapping merged exon intervals per gene, discarding exonic segments that overlap another gene’s exons on the same strand. For each gene we collect the coverage bins overlapping its exons across all test sequences and strand-matched tracks, and undo the training-time squashing transforms to recover coverage on the original scale. A gene expression estimate per track is the mean exonic coverage scaled by exon length in bins (i.e. total exonic coverage), optionally with a pseudocount equal to a fixed quantile of nonzero coverage, followed by a log_2_(*x* + 1) transform. This yields predicted and measured log-expression matrices over genes *×* tracks. Raw accuracy is the per-track Pearson r and explained-variance *R*^2^ between predicted and measured log-expression across genes.

#### Expression specificity

As for coverage, per-gene mean expression dominates raw accuracy. Within each target group we quantile-normalize the log-expression across genes to a common reference and, for each gene, subtract the mean across the group’s tracks. The per-track Pearson *r* and *R*^2^ on these normalized values measure how well relative expression differences between samples are predicted, independent of each gene’s overall level.

#### Highly variable genes

To assess accuracy per gene across samples, we correlate the quantile-normalized predicted and measured log-expression across the tracks within a group, computing a Pearson *r* and *R*^2^ per gene. Because most genes vary little across samples, we summarize over highly variable genes—the top 20% by measured log-expression standard deviation across the group.

#### Model comparison tests

Bin- and gene-level accuracy is compared across replicates with different training and test splits. The two models share the split at each replicate index, which is what the pairing exploits. Each replicate contributes one value, the mean over the tracks or genes the panel plots, and the two models are compared by a two-sided paired t-test on those values. The plotted points are per-track or per-gene accuracy pooled over replicates, so the points and the test summarize different units. Cerberus against Borzoi is the exception; the two were trained on different fold splits, so their replicates cannot be paired, and that comparison instead pairs on the track, tissue or gene the two models share and applies a two-sided Wilcoxon signed-rank test.

#### Variant-effect comparison tests

A fine-mapped QTL is called separately in each tissue it acts in, and its measurements and predictions correlate strongly across those tissues. Across the eQTL benchmark a variant is fine-mapped in 2.4 tissues on average, and the intraclass correlation across those tissues is 0.83 for the measured effect size and 0.88 to 0.89 for the models’ absolute predicted effects (Supplementary Fig. S9). The variant is therefore the unit of replication in every variant-effect comparison.

Models are compared by a cluster bootstrap over variants. Positive and negative variants are resampled separately with replacement within each stratum, which holds fixed the class ratio that AUPRC depends on; each drawn variant carries all of its tissue rows, and both models see the same resample. Then p is twice the smaller tail of 2,000 resampled differences about zero, floored at 1/2,000 (*^∗^ p* < 0.05, *^∗∗^ p* < 0.01, *^∗∗∗^ p* < 0.001; uncorrected across strata).

The violin and scatter panels score per-tissue cells: within each (tissue, stratum) cell, classification AUPRC over each positive variant and its matched negative, ranking by absolute predicted effect, and Spearman *ρ* between signed predicted and measured effect on positives. Cells with fewer than 20 positives are dropped and logged. The split violins show these per-tissue estimates, and each resample takes the median across tissues of the difference between models. The per-tissue scatter panels treat every variant as one stratum and take the mean across tissues, matching the means they annotate. The forest panels of Supplementary Fig. S11 instead collapse each variant to one row, averaging its prediction over the tissues in which it is a positive, and score each stratum once on those rows under both metrics, reporting a 95% percentile interval; strata holding fewer than 20 variants carry an interval but no test. These estimands differ because collapsing variants changes tissue weighting and pools scores into a single ranking.

Each violin half is a Gaussian kernel density estimate truncated at the observed range and normalized to equal maximum width, with interior lines at the quartiles, and every tissue is overlaid as a point.

For the sQTL and paQTL comparison of Supplementary Fig. S10, positives more than 100 kb from the nearest annotated splice or poly-A site are excluded along with their matched negatives; the benchmark sets themselves are unfiltered.

### 5.6 GTEx fine-mapped QTL benchmark

To evaluate sequence-to-function models on cis-regulatory variant effect prediction, we constructed positive and matched-negative variant sets from GTEx v11 for three molecular phenotypes—gene expression (eQTL), pre-mRNA splicing (sQTL), and alternative polyadenylation (paQTL)—across the 50 GTEx tissues. Positives are statistically fine-mapped causal variants (SuSiE posterior inclusion probability, PIP *≥* 0.9); each is matched one-to-one to a negative drawn from the same modality’s pool of nominally significant cis variants. Matching controls for local genomic context, expression of the linked gene, distance to the nearest cognate regulatory feature, minor allele frequency (MAF), and exact reference and alternate alleles. The final benchmark contains 23,836 eQTL, 20,732 sQTL, and 4,991 paQTL positive variants, deduplicated as the union across all 50 tissues. Scoring matches each GTEx tissue to that tissue’s GTEx RNA-seq tracks in our training data, which are grouped by organ and include no cultured-cell group. Thus, every model evaluation runs on the remaining 48 tissues, while the panels describing the release itself use all 50. The benchmark sets and the matching code are available in the qtl-bench repository [34].

#### 5.6.1 Data sources

We used GTEx v11 cis-QTL summary statistics and SuSiE fine-mapping from the GTEx Consortium [54]. For each modality (eQTL; sQTL from LeafCutter intron-cluster usage; paQTL from 3*^′^* UTR usage ratios) we downloaded the `signif_pairs` files (nominally significant variant–phenotype pairs from tensorQTL) and the SuSiE summary files (per-(variant, phenotype) PIP, credible-set membership, and allelic fold change) for all 50 tissues. We restricted the SuSiE summaries to phenotypes with `biotype == “protein_coding”`, dropped variants with missing median TPM in the focal tissue, and retained SNPs only (indels excluded).

Gene annotation used GENCODE v48 basic protein-coding transcripts on GRCh38, excluding readthrough transcripts. Canonical transcripts were taken from the GTF, preferring `MANE_Select` and falling back to `GENCODE_Primary` for genes lacking a MANE assignment. Expression came from the GTEx v11 RNA-SeQC v2.4.3 gene-level median-TPM matrix.

#### 5.6.2 Positive variant definition

For each tissue and modality, a positive variant–phenotype pair was any SuSiE summary row with PIP *≥* 0.9. Nominal p-values and per-allele slopes (sQTL/paQTL) were looked up from the corresponding signif pairs file for the output VCF. When a variant entered the positive set against multiple phenotypes—for example, overlapping eQTL credible sets for adjacent genes in the same cis window—we collapsed to one row per variant: eQTL used region precedence (see below), while sQTL and paQTL kept the maximum-PIP row.

#### eQTL

Each (variant, gene) positive was labeled relative to its gene:

- *3^′^ UTR* if the variant lies within any 3*^′^* UTR interval of the gene;
- *CDS* if it lies within any CDS interval;
- *5^′^ UTR/non-exonic* otherwise, meaning the variant overlaps neither a 3*^′^* UTR nor a CDS interval of the gene, and carrying tss dist, the minimum absolute distance to any TSS of the gene. Annotation is restricted to protein-coding transcripts, so the class’s only exonic sequence is 5*^′^* UTR.

When a variant overlapped a 3*^′^* UTR and a CDS interval across different transcripts, the canonical transcript determined the label; if the canonical transcript covered neither, the variant defaulted to *3^′^ UTR*. Cross-gene multi-mappings collapsed to a single row with precedence *3^′^ UTR* > *CDS* > *5^′^ UTR/non-exonic*, ties broken by smallest TSS distance.

#### sQTL

Splice sites were the internal exon boundaries of basic protein-coding transcripts—every non-terminal exon end (donor) and non-first exon start (acceptor), per transcript, deduplicated by (chrom, position, gene). Each positive carried the minimum distance to a splice site of its gene; positives whose gene had no splice site were dropped.

#### paQTL

Polyadenylation sites (PAS) were proxied by the annotated 3*^′^* end of every transcript, gene-tagged. Each positive carried the minimum distance to a PAS of its gene.

#### 5.6.3 Negative candidate pool

Per modality, we built one cross-tissue candidate pool by unioning all 50 tissues’ `signif_pairs`, deduplicating to one row per (variant, phenotype), and taking the median allele frequency across the tissues in which the pair was observed. Pooling keeps the candidate set dense in low-power tissues, which have few significant pairs of their own. For each focal tissue we restricted the pool to protein-coding genes with nonzero median TPM and at least one SuSiE row in that tissue, then excluded every variant with PIP *≥* 0.01 for any protein-coding phenotype in any tissue. This cross-tissue blocklist prevents weakly fine-mapped variants in one tissue from serving as negatives in another.

A candidate’s `signif_pairs` gene may lie far from any regulatory feature. So that the matching distance reflects the nearest plausibly regulated gene, we reassigned each negative to the gene owning the genome-wide nearest cognate feature: nearest TSS for eQTL TSS candidates, nearest splice site for sQTL, nearest PAS for paQTL; TPM was reloaded from the reassigned gene. 3*^′^* UTR and CDS eQTL negatives kept their original gene, since compartment overlap rather than distance drives their matching.

Within each tissue and modality, we paired each positive to one negative, first bucketing on exact alleles and (for eQTL) region: (region, ref, alt) for eQTL, (ref, alt) for sQTL and paQTL. Within a bucket, each candidate was scored by its dissimilarity to the positive,

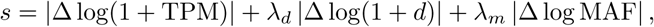

where Δ is the difference between candidate and positive, d is the modality-specific feature distance, and λ*_d_* = λ*_m_* = 1; the distance term was suppressed for 3*^′^* UTR and CDS eQTL buckets, where compartment overlap already controls for genomic context. Positives then claimed negatives greedily, each taking the unclaimed candidate in its bucket with the lowest s, so no negative was used twice.

#### 5.6.4 Variant scores

We scored each SNP–gene pair with one of two gene-restricted statistics: the log fold change (logFC) for expression QTLs (eQTLs), and the maximum profile shift (MPS) for splicing QTLs (sQTLs) and polyadenylation QTLs (paQTLs). For each SNP–gene pair, reference and alternate sequences were predicted separately, each as the average of the forward and reverse-complement passes; the reverse-complement coverage track was spatially reversed and strand-pair permuted back into forward coordinates before averaging. Targets were restricted to gene-appropriate assays and to those whose strand is compatible with the gene’s annotated strand, with unstranded targets retained for both orientations. Let 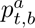 denote the predicted coverage for allele *a ∈* {ref, alt}, target *t*, and bin *b*.

The logFC sums predicted coverage over the set of output bins *E_g_* overlapping the gene’s exons, cropped to the model’s output window, and compares the alleles on a log scale,

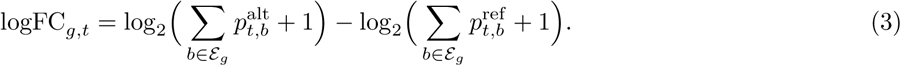

It is signed relative to the GTEx effect allele.

The MPS quantifies variant effects on the normalized shape of log-transformed predicted coverage across a gene. Predictions were restricted to the set of output bins *B_g_* spanning the gene locus, from the gene’s first exon start to its last exon end, with interval boundaries mapped to bins at the model’s output stride by rounding to the nearest bin (requiring > 50% overlap) and cropped to the model’s output window. Unlike the exon bins *E_g_* of the logFC, this span includes introns, whose coverage carries much of the splicing and intronic polyadenylation signal that sQTL and paQTL effects alter. Within the gene-restricted window, predictions were log-transformed and normalized to a discrete profile over bins,

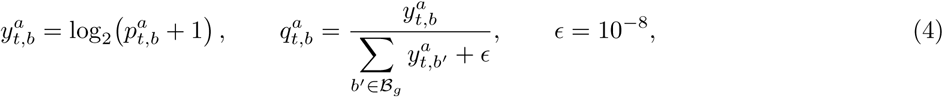

separately for each allele. The MPS is the L*_∞_* (Chebyshev) norm of the difference between these two profiles, computed per target,

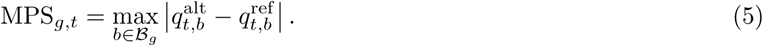

MPS reports the largest absolute per-bin difference between log-coverage profiles, with per-allele normalization intended to control for overall expression level. This control is approximate because the logarithm precedes normalization. It is unsigned and bounded in [0, 1].

#### 5.6.5 QTL classification scoring

Every variant carries exactly one designated gene—positives their fine-mapped QTL gene, negatives the gene assigned during matching—and is scored only there, so gene-dense loci accumulate no advantage over gene-sparse ones. Both models were scored with identical settings: forward and reverse-complement passes averaged, and the target set restricted to RNA-seq tracks. Each score was computed per replicate and then averaged across the eight Cerberus folds and the four Borzoi replicates. A variant’s score in GTEx tissue T is then the mean score over the GTEx RNA-seq training tracks of *T*’s tissue type—the same tracks for both models, and the only point at which tissue identity enters the prediction. Per-tissue AUROC and AUPRC rank that single scalar, in absolute value for logFC, over the tissue’s positives and their matched negatives; for eQTLs, Spearman ρ compares the signed logFC with the measured effect size over the positives.

A (variant, gene) pair is scorable only when at least one of the gene’s scored bins—its exons for logFC, its span for MPS—falls within the model’s output window centered on the variant, the central 196,608 bp of Borzoi’s 524,288 bp input after its crop and the full 786,432 bp for Cerberus; a partially covered gene is scored over its covered bins. We assign unscorable pairs a score of zero and retain them in the ranking rather than dropping them, so that each model is held responsible for the variants its receptive field cannot reach; reach is part of what the comparison measures. Unscorable pairs make up 17.7% of eQTL positives and 24.3% of their matched negatives for Borzoi, against 5.4% and 6.1% for Cerberus. The sQTL and paQTL comparison is restricted to variants within 100 kb of the cognate site, where the rates are 4.9% and 7.0% for Borzoi against 0.0% and 0.2% for Cerberus on sQTLs, and 3.8% and 4.4% against 0.4% and 0.1% on paQTLs. Scoring only the pairs both models reach leaves Cerberus ahead overall on both phenotypes (p *≤* 0.002), with the gain again concentrated near the affected site.

#### 5.6.6 AlphaGenome scoring

AlphaGenome’s non-commercial license does not permit us to run it outside the API, so we scored it against a self-deployed Vertex AI Model Garden endpoint, calling the public `alphagenome` SDK (v0.4.2.8) through the `alphagenome_cloud_sdk` transport (v0.4.2.10). The endpoint serves the released distilled student, which its authors trained against a 64-model all-folds teacher ensemble [2].

We scored each variant with one score variant call, passing the variant, a 1 Mb interval centered on it, and AlphaGenome’s recommended RNA-seq variant scorer RNA SEQ. That scorer reports the log ratio of predicted expression between the alternate and reference alleles under a gene’s exon mask, the statistic we compute for Cerberus and Borzoi. Each call returns one signed score per variant, gene, and GTEx tissue. A variant’s AlphaGenome score in tissue *T* is the mean of these scores over the GTEx tissues of *T*’s tissue type, the grouping applied to Cerberus and Borzoi tracks.

The API takes a genomic interval and a variant and exposes no control of sequence orientation, so AlphaGenome scores rest on the forward strand alone, whereas Cerberus and Borzoi are scored as the average of a forward and a reverse-complement pass. Its authors applied that same strand-averaging augmentation in their own variant evaluations and report that it “did not consistently enhance the performance of AlphaGenome’s distilled model evaluations” [2].

We did not compare AlphaGenome on the sQTL and paQTL benchmarks, whose API scorers measure log fold change across a gene’s annotated splice junctions or between its annotated proximal and distal cleavage sites, rather than over predicted coverage as the maximum profile shift does, confounding a difference between models with a difference between statistics. Those scorers also returned a score at the designated gene for 78% of sQTL and 52% of paQTL variants within 100 kb of the cognate site, against 99.8% for Cerberus on both, a gap we could not attribute to a documented property of the scorers.

### 5.7 Hydra block interpretation

We extracted the parameters and internal activations of every Hydra block of each of the eight Cerberus replicates, on 16 sequences drawn at random from that replicate’s test split. Each block contains 20 heads, and each head has a scalar decay parameter a and an input-dependent step size Δ*_t_* at each position *t*. The input and readout factors *b_t_* and *c_t_* have shape (*n*_pos_, *n*_groups_, *d*_state_); because *n*_groups_ = *d*_state_ = 1, they reduce here to vectors over positions and are shared across the 20 heads of a block. The forward and reverse recurrences share a; we extracted Δ*_t_*, b*_t_* and c*_t_* separately for each, together with the mean absolute output of each pass, which measures how much each direction contributes.

All statistics below drop the outermost 256 bins (8 kb) at each end of the 786 kb window. The recurrence warms from zero at each window edge, spiking Δ to 6–7 times the interior mean over the outermost 10 bins of the seventh block, an artifact of the boundary rather than the sequence. Every reported quantity changes little as the trim varies from 64 to 2,048 bins.

#### Reach

For *i* > *j*, the Mamba-2 mixing weight can be written as *m_ij_* = *c_i_* exp(*s_i_−s_j_*)*b_j_*Δ*_j_*, where *s_t_* = *aΣ_k≤t_* Δ*_k_* and the decay parameter a < 0; the reverse pass supplies *i* < *j* in the same form. The factor exp(*s_i_* − *s_j_*) controls how information from position j decays with distance. For each head and pass, we take the median step size Δ̃ over positions, averaged over windows, and define an effective half-life of ln 2/(*|*a*|*Δ̃) bins, the distance over which exp(*s_i_* − *s_j_*) falls by half at constant step size. We use the median because Δ is heavy-tailed: in a typical seventh-block head the top 1% of positions carry 40% of the summed Δ, so the mean would reflect the spikes rather than the decay between them. A head’s half-life is the harmonic mean of its forward and reverse half-lives, equivalently the half-life at the mean of the two passes’ median step sizes. The ratio of forward to reverse half-life compares the two directions. We report half-lives in kilobases at 32 bp per bin and summarize their distribution within a block by quartiles. For every half-life threshold between 8 and 128 kb, the fraction of heads above it rises from the first block to the seventh.

#### Operator coarse-graining

A head’s operator *m_ij_*, summed over the two passes, is 24,576 *×* 24,576 per window, too large to display. We coarse-grained it onto a 512 *×* 512 grid of 1,536 bp cells, cell (*I*, *J*) holding mean*_i∈I_Σ_j∈J_ m_ij_*, the total weight an output position in *I* draws from the sources in *J*. Because *b* and *c* are signed, cells can be negative, so we use their absolute values, after dropping 6 cells at each edge to cover the 256-bin trim. On-diagonal weight is the share of a head’s total weight within two cells of the diagonal, and column weight is the total that the sources in a column write into the window.

#### Anchors and enrichment

A head’s anchors in a window are the top 1% of its trimmed positions by step size averaged over the two passes, Δ = (Δ_fw_ + Δ_rev_)/2. We intersected them with GENCODE v48 transcription start sites, exons, introns and UTRs, and with ENCODE cCREs and CTCF ChIP-seq peaks, binned to the same 32 bp grid. Fold enrichment is the number of anchors overlapping a class divided by the number expected from the fraction of scored bins the class covers. Significance comes from a circular-shift null. Rolling the anchor mask along the window preserves both the anchor count and the local clustering of anchors while destroying their alignment to the annotation. The circular cross-correlation of mask and annotation gives the overlap at all L shifts from a single pair of FFTs, so the p-value is an exact rank among the admissible shifts rather than a normal approximation. Shifts of fewer than 1,000 bins (32 kb) in either direction are excluded, because anchors cluster locally and a small roll leaves them correlated with the annotation. Anchor masks are pooled over the heads and windows of a block, giving one test per block and replicate rather than one per head. Heads within a deep block anchor at overlapping positions (median pairwise Jaccard 0.11 in the seventh block, against 0.004 for the shifted null), so 20 heads are not 20 independent tests, whereas the eight replicates are independently trained models. Two splits control for confounding, under the same null and pooling. To separate promoter identity from GC content, promoter and non-promoter bins are each split at the median GC content of promoter-like cCREs. To separate CTCF from promoter signal, CTCF ChIP-seq peaks are split on overlap with a promoter-like cCRE. Separately, a GC-matched expectation computes each anchor’s expected overlap only from bins in its own GC decile.

#### Anchor placement

For each annotation class we took the midpoints of its features from the BED file, dropped those within 2 kb of a trimmed window edge, and averaged the fraction of anchors at each signed offset out to *±*2 kb in 32 bp bins, divided by its expectation under uniform placement.

### 5.8 Manuscript preparation

Claude (Anthropic) was used to assist with writing and editing of this manuscript. All scientific content, interpretations, and conclusions are the authors’ own. All AI-assisted text was reviewed and revised by the authors, who take full responsibility for the accuracy and integrity of the work.

## Abbreviations

AUPRC: area under the precision–recall curve
AUROC: area under the receiver operating characteristic curve
CAGE: cap analysis of gene expression
cCRE: candidate cis-regulatory element
CDS: coding sequence
eQTL: expression quantitative trait locus
ISM: in-silico mutagenesis
MAF: minor allele frequency
MPS: maximum profile shift
paQTL: polyadenylation quantitative trait locus
PAS: polyadenylation site
PIP: posterior inclusion probability
QTL: quantitative trait locus
RoPE: rotary positional embedding
SNP: single-nucleotide polymorphism
sQTL: splicing quantitative trait locus
SSD: state space duality
SSM: state space model
TPM: transcripts per million
TSS: transcription start site
UTR: untranslated region.

## Declarations

### Ethics approval and consent to participate

Not applicable. This study is computational and analyzed only previously published, publicly available human and mouse data.

### Consent for publication

Not applicable. This study contains no data from any individual person.

### Availability of data and materials

The models, training code, and evaluation code described in this article are available in the Baskerville repository [29] under the Apache License 2.0, which also links the training data. The GTEx v11 cis-QTL summary statistics and SuSiE fine-mapping results analyzed in this study are available from the GTEx portal [54]. The GTEx v11 fine-mapped eQTL, sQTL, and paQTL benchmark sets constructed here, comprising positive variants and their matched negatives, and the code that constructs them are available in the qtl-bench repository [34] under the Apache License 2.0. The full inventory of training tracks and their sources is included as Supplementary Table S1.

### Competing interests

DK, HY, and XH are employees of Calico Life Sciences LLC. JL consults for Calico Life Sciences LLC.

### Funding

This work was funded by Calico Life Sciences LLC.

### Authors’ contributions

DK conceived the study, designed and performed the block-comparison experiments, trained and evaluated the models, developed the fine-mapped QTL benchmarking framework, and wrote the manuscript. HY developed the analyses interpreting the learned Hydra parameters, in iterative discussion with DK. XH analyzed candidate eQTL examples across Cerberus, Borzoi, and AlphaGenome and developed the example selection analysis. JL contributed to the conception of the study and advised on model training and evaluation throughout. All authors read and approved the final manuscript.

## Acknowledgements

We thank Ian Holmes for implementing and benchmarking efficient sequence-mixing architectures, including Mamba, in JAX during his sabbatical at Calico Life Sciences, which motivated our interest in state space models for regulatory sequence modeling [55]. We thank Ben Auerbach, Divyanshi Srivastava, and David Wang for discussions of state space models in group meetings and journal clubs.

## Supplementary information

Supplementary Figures S1–S15 and Supplementary Table S1 accompany this preprint. Table S1 gives the complete set of human and mouse coverage tracks used to train Cerberus on separate sheets of 8,361 and 3,102 tracks, one row per track, with assay, source study or consortium, sample or cell type, accession, and the transformation applied during dataset construction, preceded by a summary sheet of track counts by source and assay. The figures and their legends follow.

## Supplementary figures

**Fig. S1.**
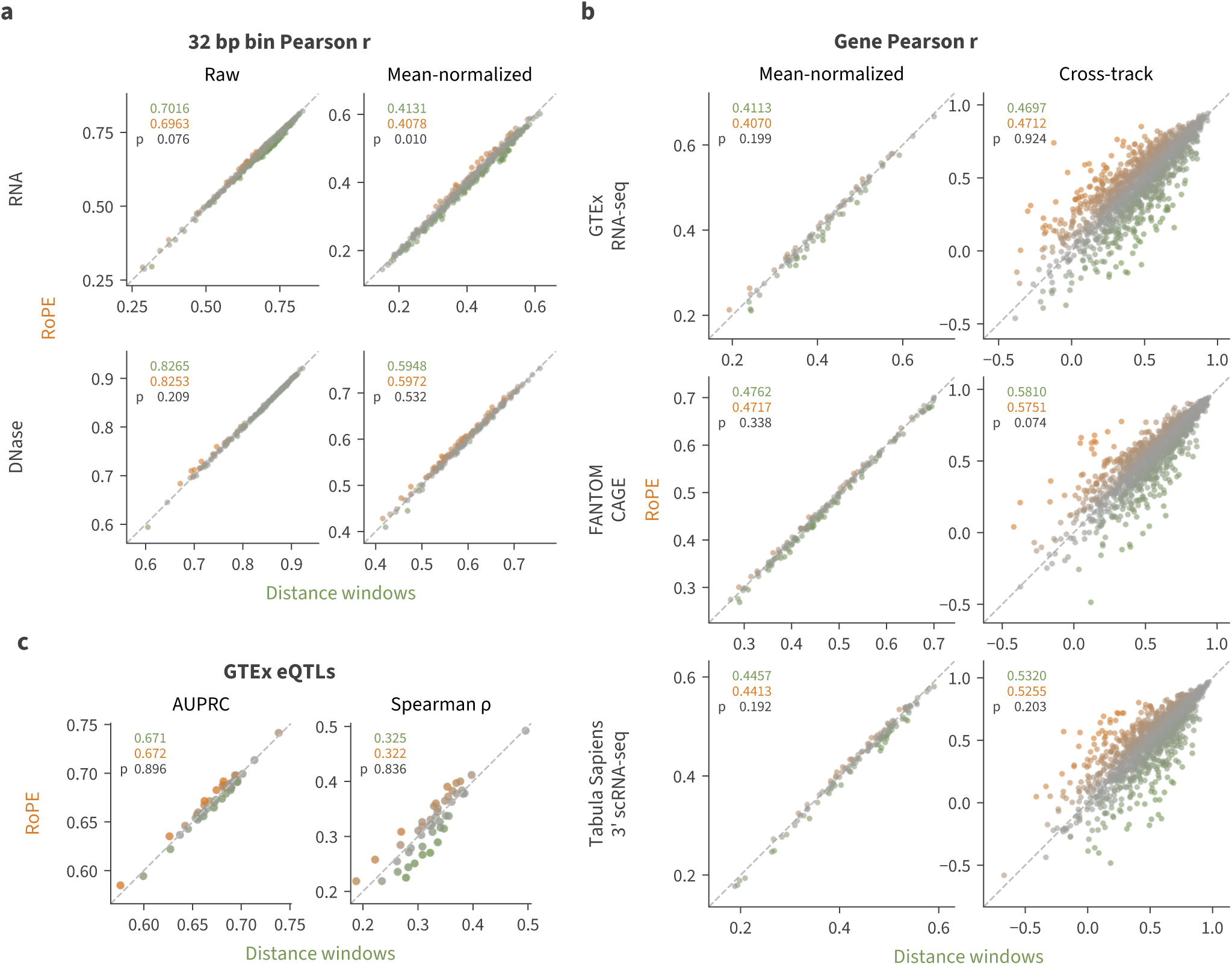
RoPE versus log-spaced distance windows. **(a)** Bin-level metrics, averaged across four replicates’ unique test sets. First row RNA tracks, second row DNase tracks. First column Pearson r across all bins for the raw values, second column Pearson r across all bins for the values after quantile normalizing each track’s genome-wide values and subtracting each bin’s mean across tracks. Points represent tracks. Means over the plotted points, with a two-sided paired t-test p across the four replicates, written in upper left (Methods). **(b)** Gene-level metrics, aggregating bin values overlapping gene exons over four replicates’ unique test sets. First row GTEx RNA-seq, second row FANTOM CAGE, third row Tabula Sapiens 3*^′^* scRNA-seq. First column Pearson r across all genes after quantile normalizing each track’s transcriptome-wide values and subtracting each gene’s mean across tracks; second column Pearson r across tracks after the same normalization, filtering to the top 20% most variable genes. In column one, points represent tracks, averaged over the four replicates; in column two, points represent genes, each held out by one replicate. Means over the plotted points, with a two-sided paired t-test p across the four replicates, written in upper left (Methods). **(c)** On the left, Classification AUPRC for GTEx fine-mapped eQTLs versus negative variants. Points represent GTEx tissues. On the right, GTEx fine-mapped eQTL coefficient Spearman ρ. Points represent GTEx tissues. Means over tissues, with the cluster bootstrap p over variants, written in upper left (Methods).

**Fig. S2.**
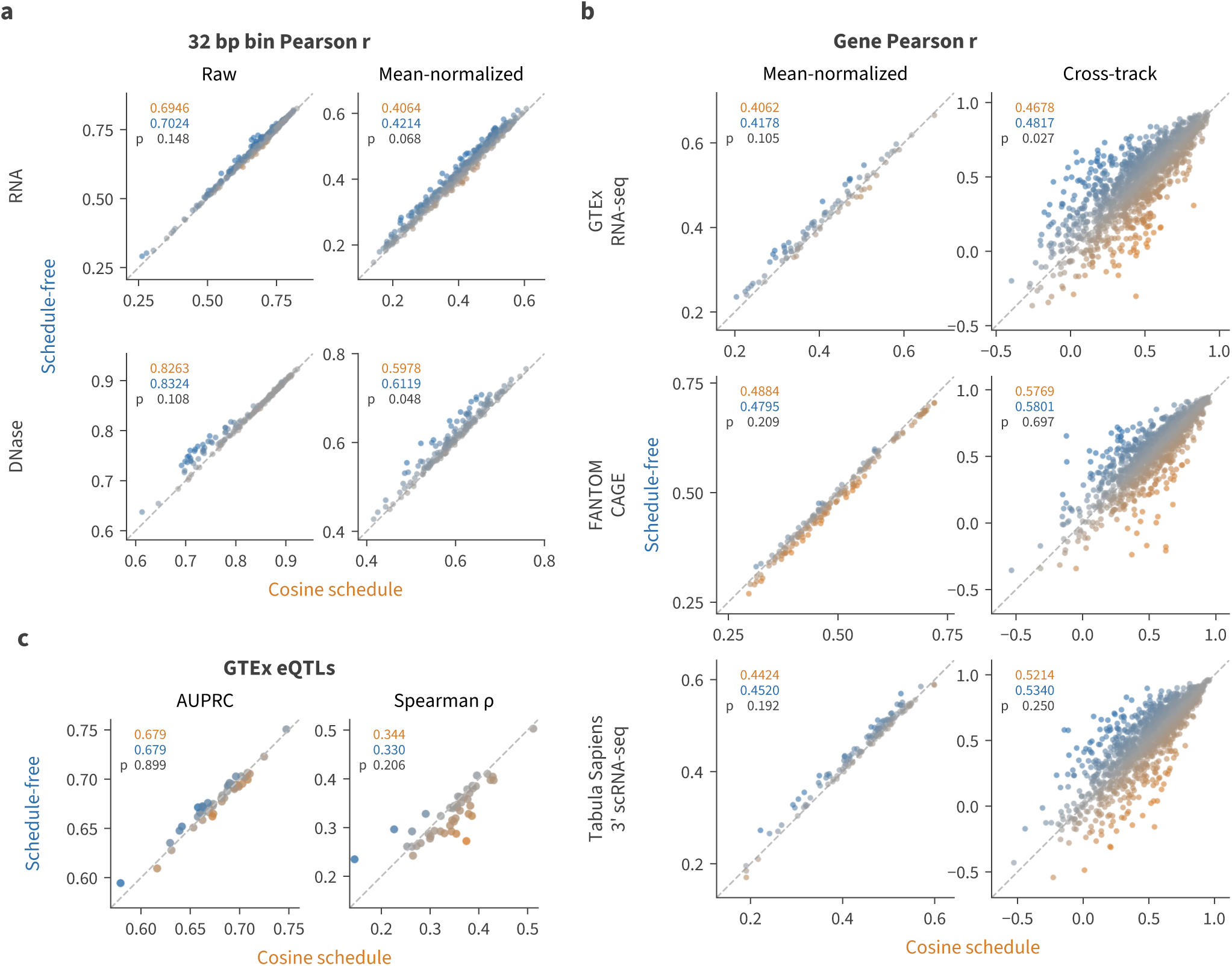
Schedule-free versus cosine decay optimization of Hydra. **(a)** Bin-level metrics, averaged across four replicates’ unique test sets. First row RNA tracks, second row DNase tracks. First column Pearson r across all bins for the raw values, second column Pearson r across all bins for the values after quantile normalizing each track’s genome-wide values and subtracting each bin’s mean across tracks. Points represent tracks. Means over the plotted points, with a two-sided paired t-test p across the four replicates, written in upper left (Methods). **(b)** Gene-level metrics, aggregating bin values overlapping gene exons over four replicates’ unique test sets. First row GTEx RNA-seq, second row FANTOM CAGE, third row Tabula Sapiens 3*^′^* scRNA-seq. First column Pearson r across all genes after quantile normalizing each track’s transcriptome-wide values and subtracting each gene’s mean across tracks; second column Pearson r across tracks after the same normalization, filtering to the top 20% most variable genes. In column one, points represent tracks, averaged over the four replicates; in column two, points represent genes, each held out by one replicate. Means over the plotted points, with a two-sided paired t-test p across the four replicates, written in upper left (Methods). **(c)** On the left, Classification AUPRC for GTEx fine-mapped eQTLs versus negative variants. Points represent GTEx tissues. On the right, GTEx fine-mapped eQTL coefficient Spearman ρ. Points represent GTEx tissues. Means over tissues, with the cluster bootstrap p over variants, written in upper left (Methods).

**Fig. S3.**
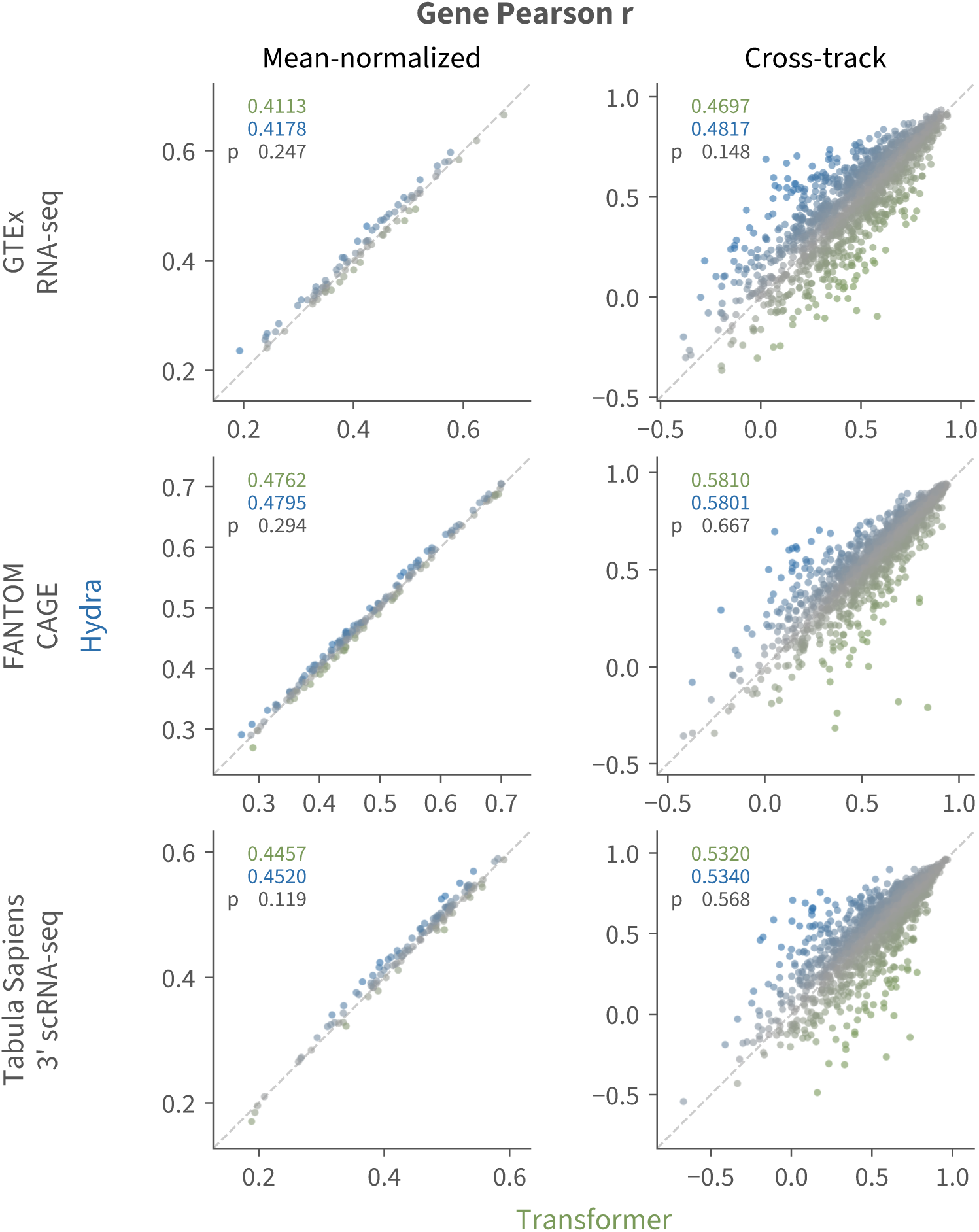
Hydra versus transformer gene accuracy. Gene-level metrics, aggregating bin values overlapping gene exons over four replicates’ unique test sets. First row GTEx RNA-seq, second row FANTOM CAGE, third row Tabula Sapiens 3*^′^* scRNA-seq. First column Pearson r across all genes after quantile normalizing each track’s transcriptome-wide values and subtracting each gene’s mean across tracks; second column Pearson r across tracks after the same normalization, filtering to the top 20% most variable genes. In column one, points represent tracks, averaged over the four replicates; in column two, points represent genes, each held out by one replicate. Means over the plotted points, with a two-sided paired t-test p across the four replicates, written in upper left (Methods).

**Fig. S4.**
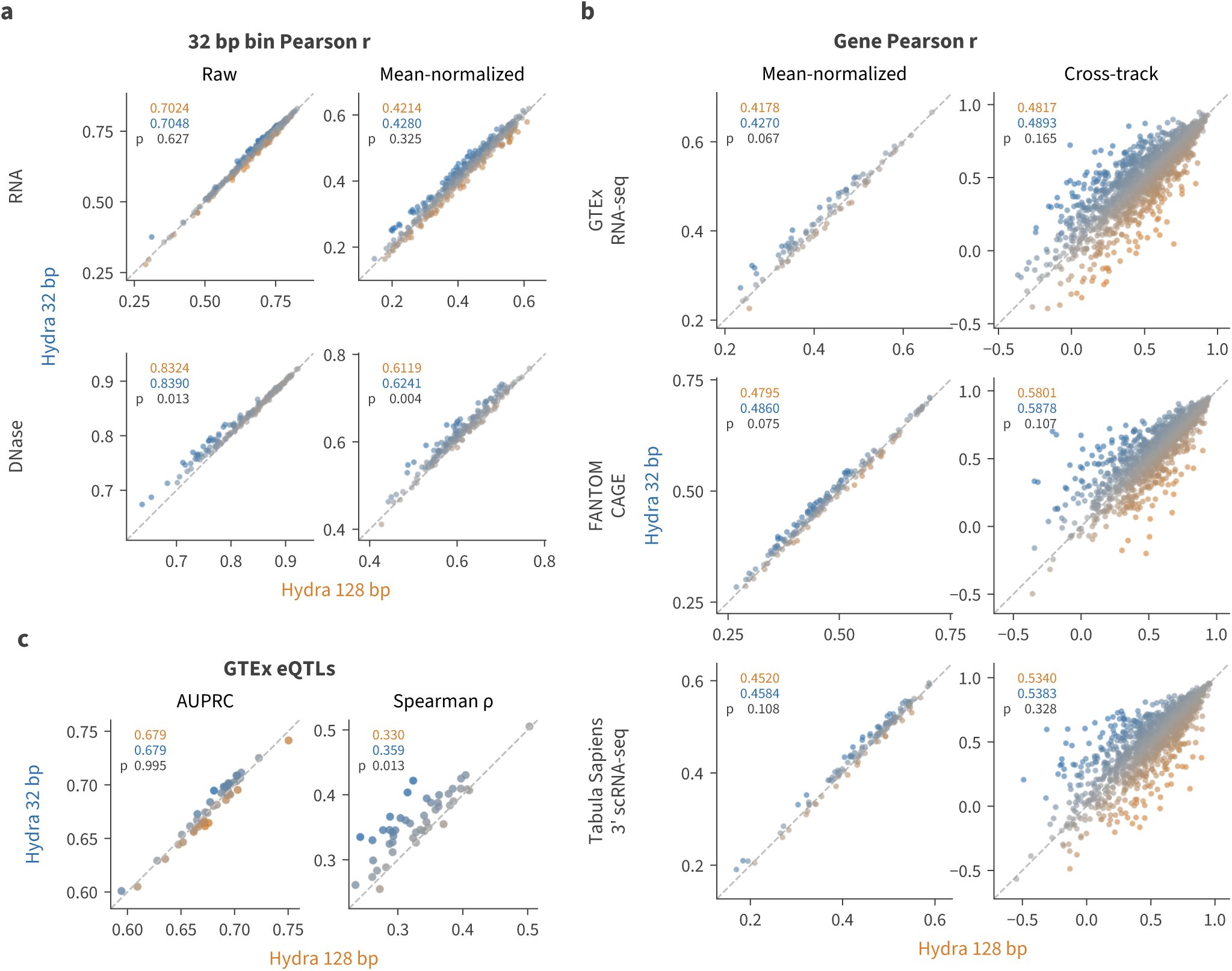
Hydra mixing at 32 bp versus 128 bp. The 32 bp model pools to 32 bp and mixes there; the 128 bp model pools to 128 bp, mixes there, and upsamples to 32 bp with a U-net decoder, which the 32 bp model does not carry. **(a)** Bin-level metrics, averaged across four replicates’ unique test sets. First row RNA tracks, second row DNase tracks. First column Pearson r across all bins for the raw values, second column Pearson r across all bins for the values after quantile normalizing each track’s genome-wide values and subtracting each bin’s mean across tracks. Points represent tracks. Means over the plotted points, with a two-sided paired t-test p across the four replicates, written in upper left (Methods). **(b)** Gene-level metrics, aggregating bin values overlapping gene exons over four replicates’ unique test sets. First row GTEx RNA-seq, second row FANTOM CAGE, third row Tabula Sapiens 3*^′^* scRNA-seq. First column Pearson r across all genes after quantile normalizing each track’s transcriptome-wide values and subtracting each gene’s mean across tracks; second column Pearson r across tracks after the same normalization, filtering to the top 20% most variable genes. In column one, points represent tracks, averaged over the four replicates; in column two, points represent genes, each held out by one replicate. Means over the plotted points, with a two-sided paired t-test p across the four replicates, written in upper left (Methods). **(c)** On the left, Classification AUPRC for GTEx fine-mapped eQTLs versus negative variants. Points represent GTEx tissues. On the right, GTEx fine-mapped eQTL coefficient Spearman ρ. Points represent GTEx tissues. Means over tissues, with the cluster bootstrap p over variants, written in upper left (Methods).

**Fig. S5.**
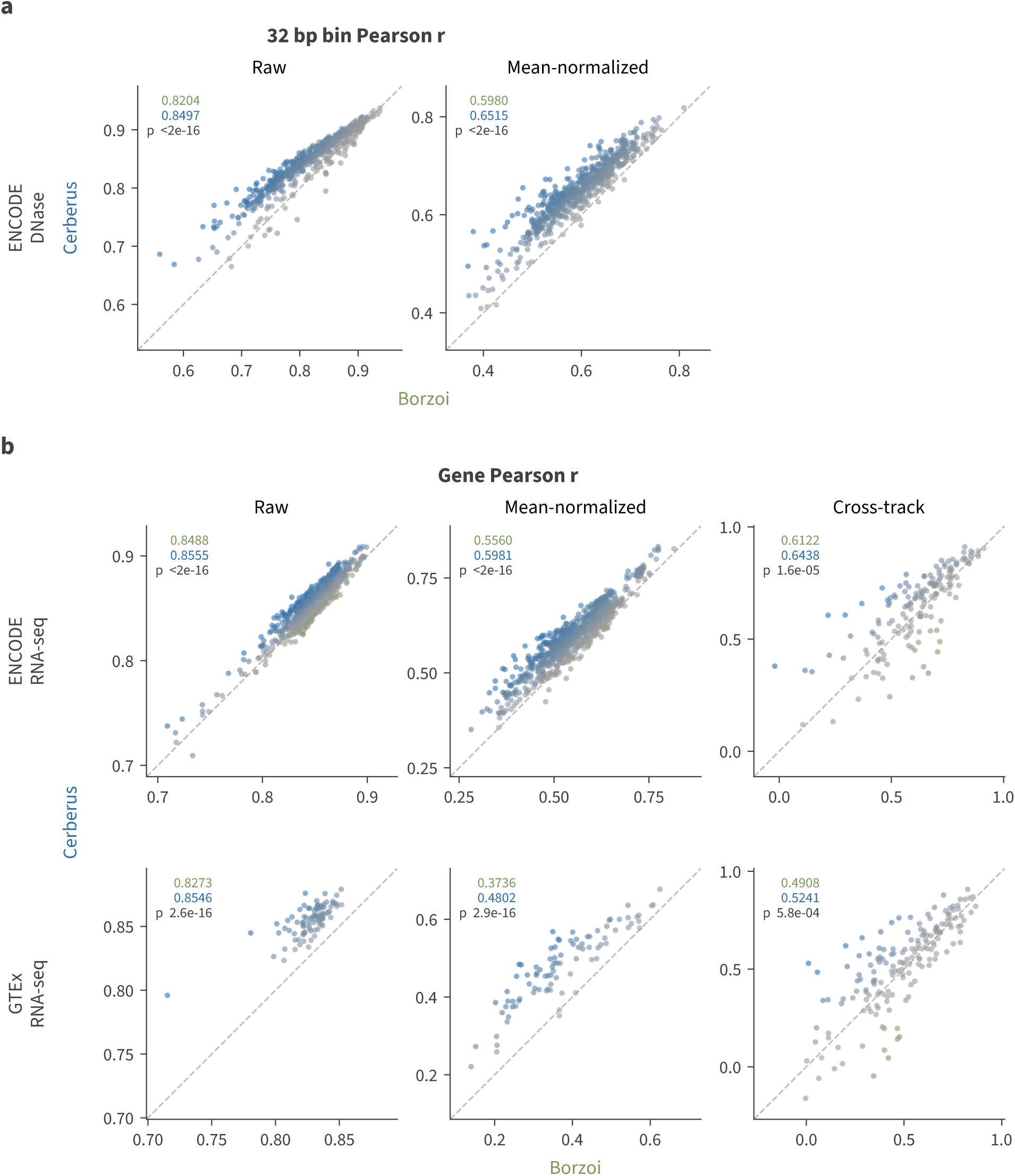
Cerberus versus Borzoi on matched tracks and genes. **(a)** Raw (left) and mean-normalized (right) per-track Pearson r for ENCODE DNase, each point a track matched between models. Other assays involved aligned read coverage transformation differences that challenge bin-level comparisons. **(b)** For gene expression, per-tissue Pearson r across genes (raw, left), the same after mean-normalization that isolates cross-tissue signal (middle), and per-gene Pearson r across tissues measuring specificity over the top 20% most variable genes (right), for ENCODE RNA-seq (top) and GTEx (bottom). Throughout, points are tinted toward the better model, the dashed line is parity, and each panel is annotated with each model’s mean over the plotted points and a two-sided paired Wilcoxon signed-rank p across them (Methods). The two models hold out different sequences and genes, so the DNase tracks in (a) rest on different test sequences and are not exactly comparable. Panel (b) avoids this by using only the 848 genes held out by both models, the intersection of Borzoi’s test genes with the union of the eight Cerberus replicates’ test genes.

**Fig. S6.**
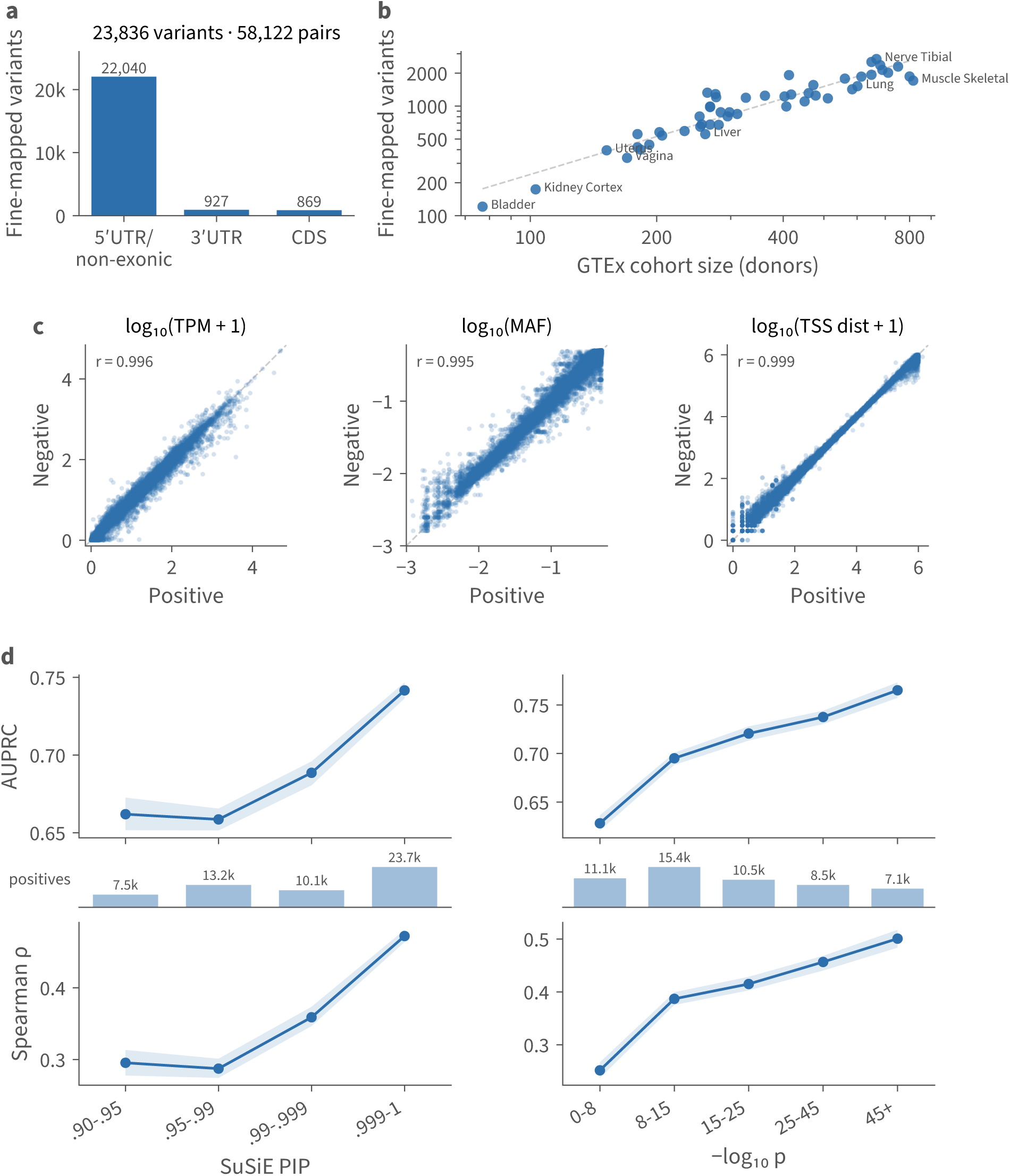
GTEx v11 fine-mapped eQTL benchmark. Positive variants are SuSiE fine-mapped causal eQTLs (posterior inclusion probability *≥* 0.9) from GTEx v11; each is paired one-to-one with a negative drawn from a cross-tissue pool of nominally significant cis variants, matched on gene-relative region and exact ref/alt alleles, linked-gene expression, minor allele frequency, and, for 5*^′^* UTR/non-exonic variants, distance to the nearest TSS. **(a)** Per-variant composition by gene-relative region. **(b)** Fine-mapped variants recovered per tissue versus GTEx donor cohort size (log–log; cohort size from allele counts); dashed line, power-law fit. **(c)** Per-pair property matching: each positive (x) against its matched negative (y) for log_10_(TPM+1), log_10_(MAF), and log_10_(TSS distance+1) (5*^′^* UTR/non-exonic variants only). **(d)** Cerberus prediction accuracy over the 48 scored GTEx tissues, stratified by SuSiE PIP (left) and association significance, *−* log_10_ p (right). Top: classification AUPRC discriminating each positive from its matched negative. Bottom: concordance (Spearman ρ) between the signed prediction and measured allelic fold change on positives. The strip between shows the number of positives per bin. Bands are 5th–95th bootstrap percentiles (200 resamples).

**Fig. S7.**
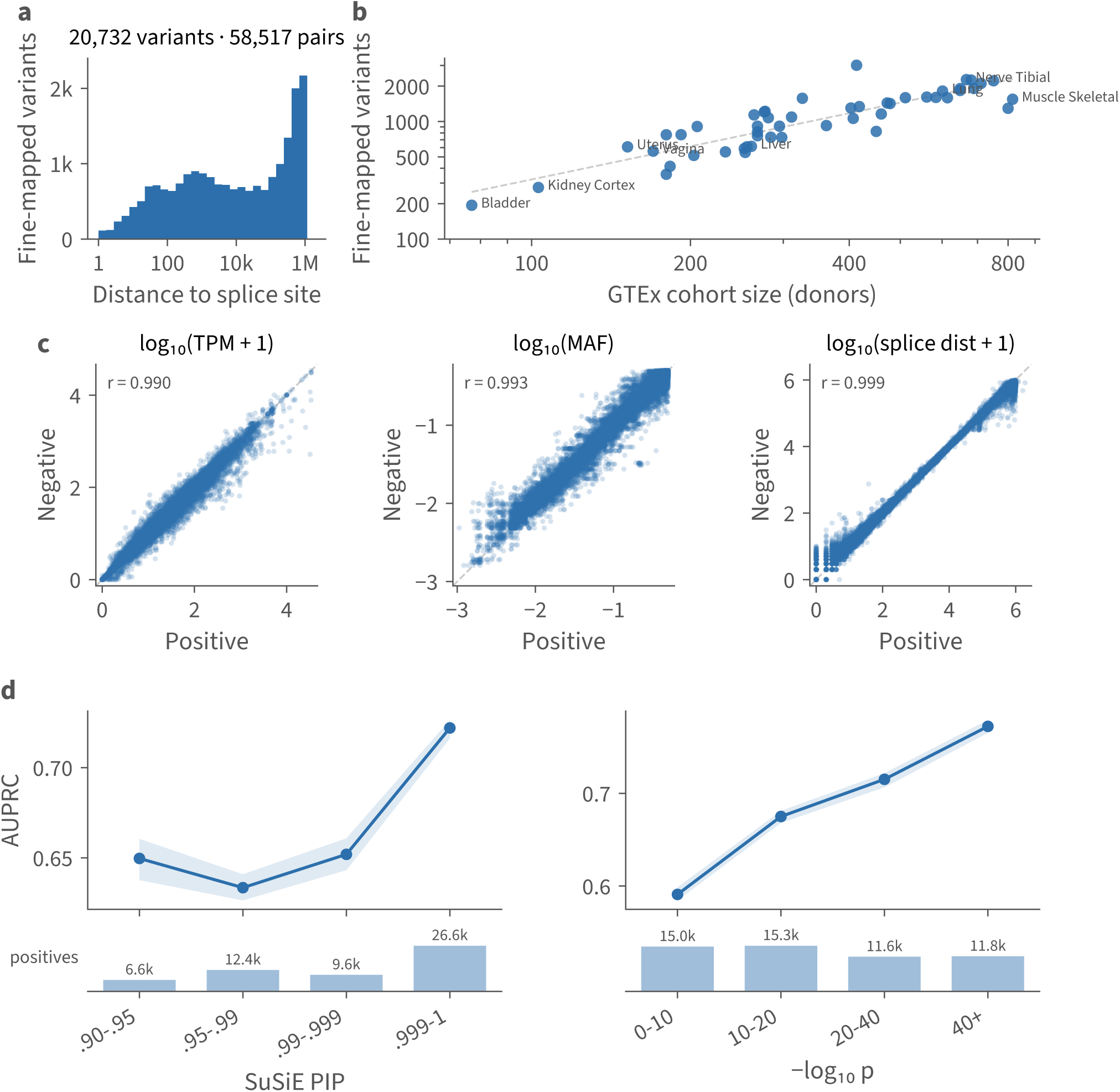
GTEx v11 fine-mapped sQTL benchmark. Positive variants are SuSiE fine-mapped causal sQTLs (posterior inclusion probability *≥* 0.9) from GTEx v11; each is paired one-to-one with a negative drawn from a cross-tissue pool of nominally significant cis variants, matched on exact ref/alt alleles, linked-gene expression, distance to the nearest splice site, and minor allele frequency. **(a)** Distance to the nearest splice site (bimodal: a splice-proximal mode and a long-range regulatory tail). **(b)** Fine-mapped variants recovered per tissue versus GTEx donor cohort size (log–log; cohort size from allele counts); dashed line, power-law fit. **(c)** Per-pair property matching: each positive (x) against its matched negative (y) for log_10_(TPM + 1), log_10_(MAF), and log_10_(splice distance + 1). **(d)** Cerberus prediction accuracy over the 48 scored GTEx tissues, stratified by SuSiE PIP (left) and association significance, *−* log_10_ p (right), by classification AUPRC discriminating each positive from its matched negative; the strip below shows the number of positives per bin. Bands are 5th–95th bootstrap percentiles (200 resamples).

**Fig. S8.**
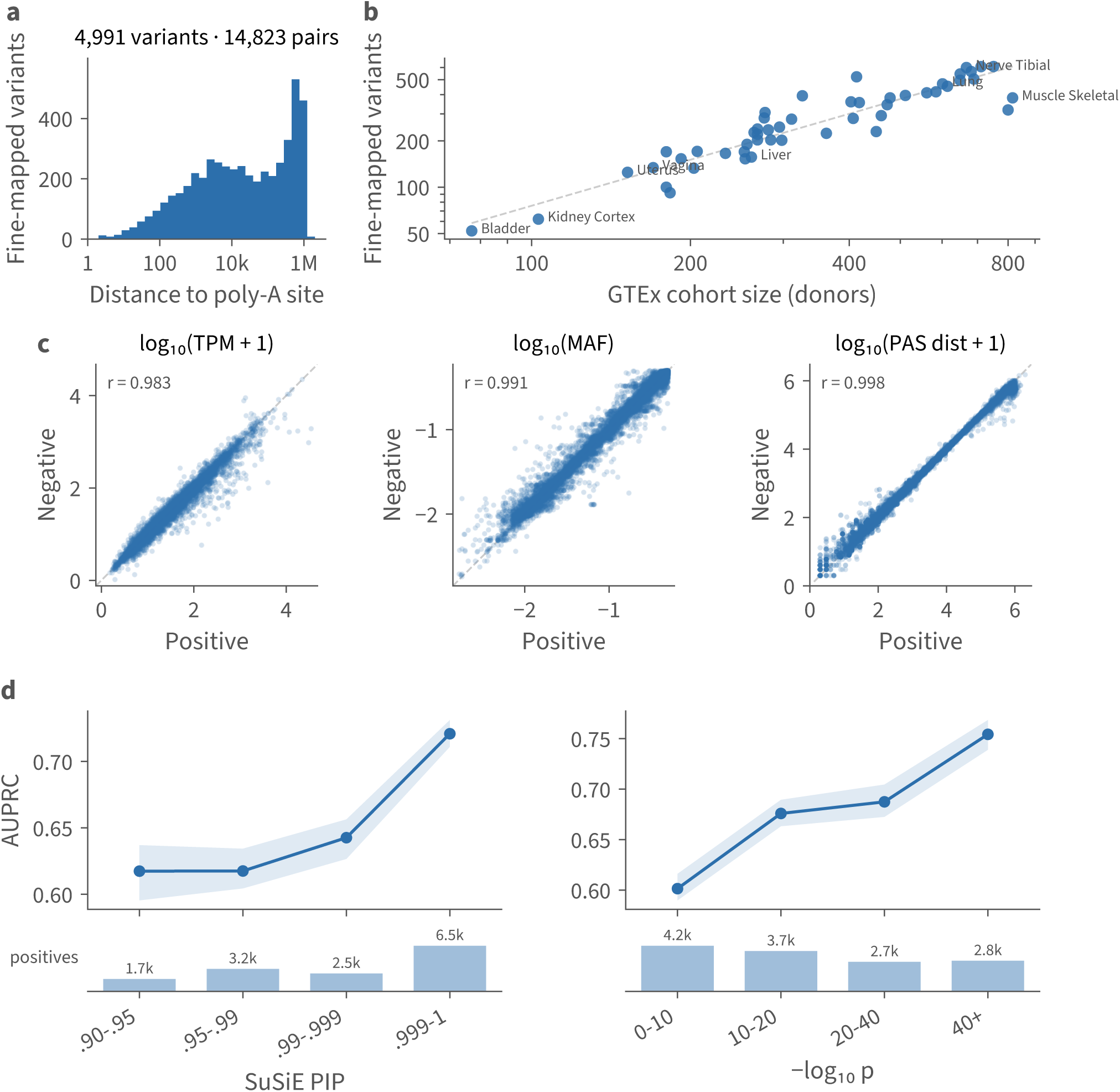
GTEx v11 fine-mapped paQTL benchmark. Positive variants are SuSiE fine-mapped causal paQTLs (posterior inclusion probability *≥* 0.9) from GTEx v11; each is paired one-to-one with a negative drawn from a cross-tissue pool of nominally significant cis variants, matched on exact ref/alt alleles, linked-gene expression, distance to the nearest poly-A site, and minor allele frequency. **(a)** Distance to the nearest poly-A site (skewed toward the long-range tail, with a strong distal mode). **(b)** Fine-mapped variants recovered per tissue versus GTEx donor cohort size (log–log; cohort size from allele counts); dashed line, power-law fit. **(c)** Per-pair property matching: each positive (x) against its matched negative (y) for log_10_(TPM+1), log_10_(MAF), and log_10_(PAS distance+1). **(d)** Cerberus prediction accuracy over the 48 scored GTEx tissues, stratified by SuSiE PIP (left) and association significance, *−* log_10_ p (right), by classification AUPRC discriminating each positive from its matched negative; the strip below shows the number of positives per bin. Bands are 5th–95th bootstrap percentiles (200 resamples).

**Fig. S9.**
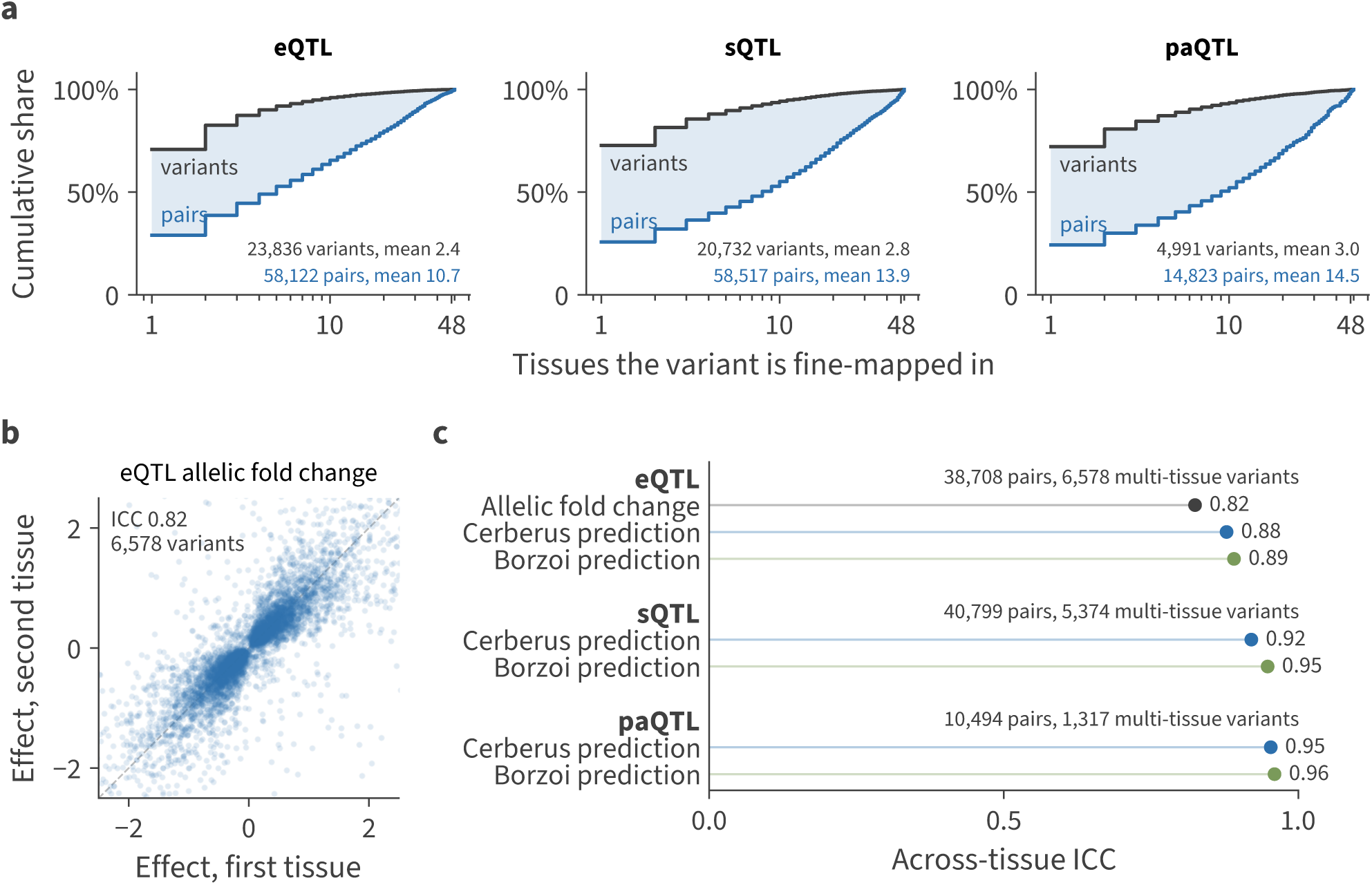
Cross-tissue structure of the fine-mapped xQTL benchmark sets. A fine-mapped QTL is called separately in every tissue it acts in, so each set is indexed by variant–tissue pair and its pairs are clustered on the variant. **(a)** Cumulative share of variants, and of variant–tissue pairs, against the number of tissues a variant is fine-mapped in. In every set the 71 to 73% of variants fine-mapped in one tissue carry 24 to 29% of the pairs, and the shaded band represents that gap. A randomly drawn variant is fine-mapped in 2.4 to 3.0 tissues, while the variant behind a randomly drawn pair is fine-mapped in 10.7 to 14.5, since a variant contributes one pair for every tissue it is found in. **(b)** Measured allelic fold change of each eQTL in two randomly chosen tissues of those it is fine-mapped in; dashed line, parity. **(c)** Across-tissue intraclass correlation of the measured allelic fold change and of each model’s absolute predicted effect, with the variant as group, over the positives scored in the 48 tissues both models cover. Splicing and polyadenylation have no allelic-fold-change analog, so the measured row is eQTL-only. Single-tissue variants carry no within-variant contrast and are excluded, so the annotated counts are the estimate’s own rather than the set’s. Because a variant’s measurements and predictions correlate strongly across tissues, variant-effect comparisons use a cluster bootstrap over variants, each resampled with all of its tissues (Methods).

**Fig. S10.**
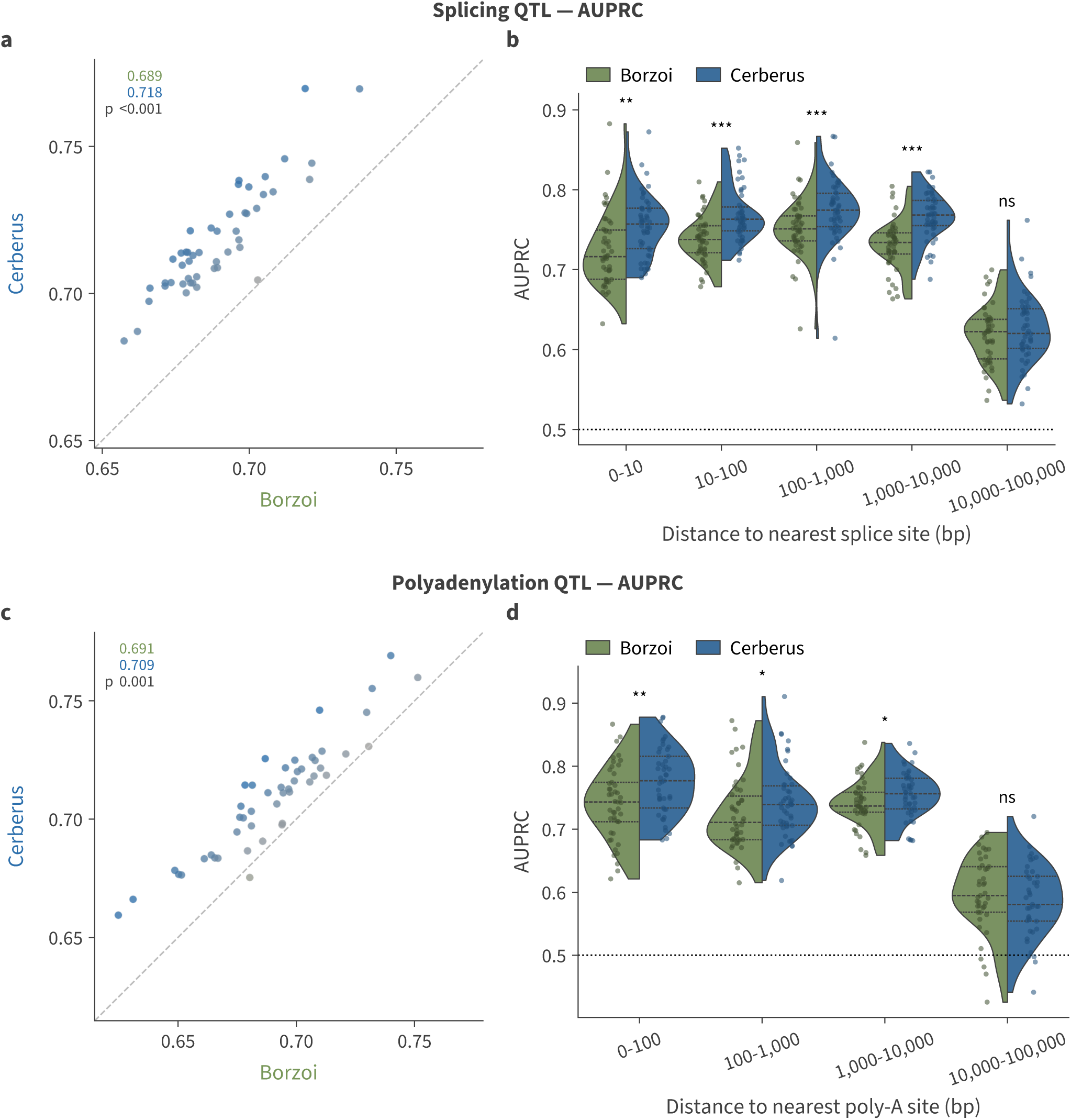
Splicing and polyadenylation QTL classification, Cerberus versus Borzoi. Fine-mapped GTEx v11 sQTLs (top row) and paQTLs (bottom row) with SuSiE PIP *≥* 0.9 in 48 tissues, each positive paired with a matched negative variant. Positives more than 100 kb from the nearest annotated splice or poly-A site are excluded, since neither model beats chance there (median AUPRC 0.49 to 0.51, against 0.50 by chance). **(a)** Per-tissue AUPRC on sQTLs; each point is one tissue tinted toward the model leading in it, the dashed line is parity, and inset values are means across tissues with the cluster bootstrap p over variants. **(b)** sQTL AUPRC stratified by the distance from the variant to the nearest annotated splice site. **(c, d)** As in (a, b) for paQTLs, by distance to the nearest annotated poly-A site. Split violins show per-tissue estimates with every tissue overlaid as a point, and stars mark a cluster bootstrap over variants (*^∗^* p < 0.05, *^∗∗^* p < 0.01, *^∗∗∗^* p < 0.001, ns otherwise); a tissue enters a bin at 20 positives (Methods).

**Fig. S11.**
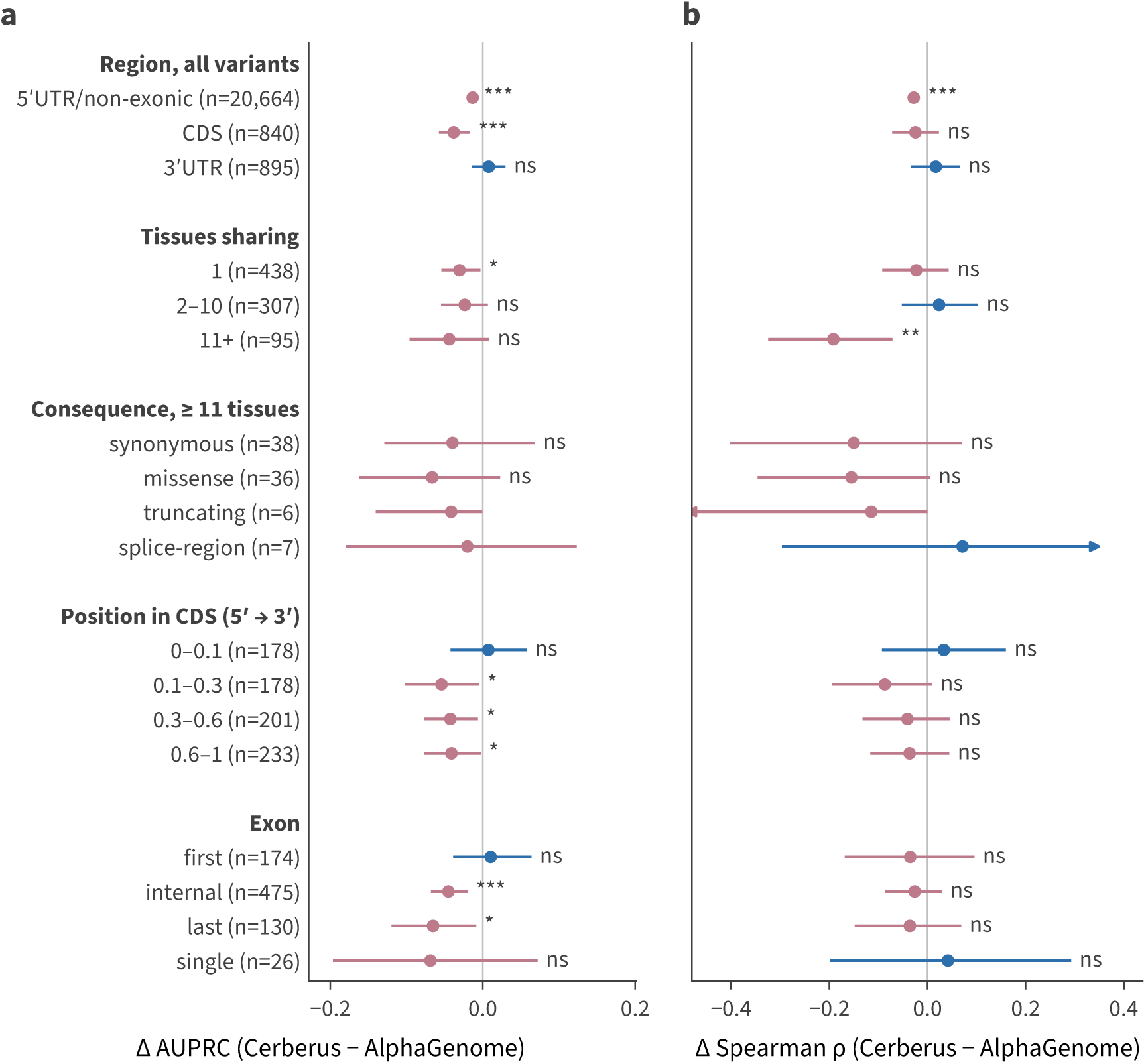
AlphaGenome’s coding eQTL advantage, stratified at the variant level. CDS-overlapping positives from the Fig. 3 benchmark (840 variants in 3,743 variant–tissue pairs) and their matched negatives, which inherit their positive’s annotation so that every stratum is a paired comparison. Each variant is collapsed to one row, averaging its prediction over the tissues in which it is a positive (Methods), so these values are not comparable with the per-tissue values of Fig. 3. **(a)** ΔAUPRC, separating positives from matched negatives, and **(b)** ΔSpearman ρ between signed prediction and measured effect over positives, under the same five stratifications: region; then within CDS, the number of the 48 tissues in which the variant is a CDS positive; snpEff consequence class among variants positive in 11 or more tissues; position along the coding sequence, as a fraction of CDS length from the start codon; and position within the exon structure of the canonical transcript. Points are the difference between models and bars 95% bootstrap intervals, with arrowheads where an interval is clipped at the axis; row labels give the number of variants, and strata below 20 carry no test (*^∗^* p < 0.05, *^∗∗^* p < 0.01, *^∗∗∗^* p < 0.001). Averaging over more tissues reduces a variant’s noise, so the tissue-sharing strata were recomputed from one randomly chosen tissue per variant, and no verdict changes.

**Fig. S12.**
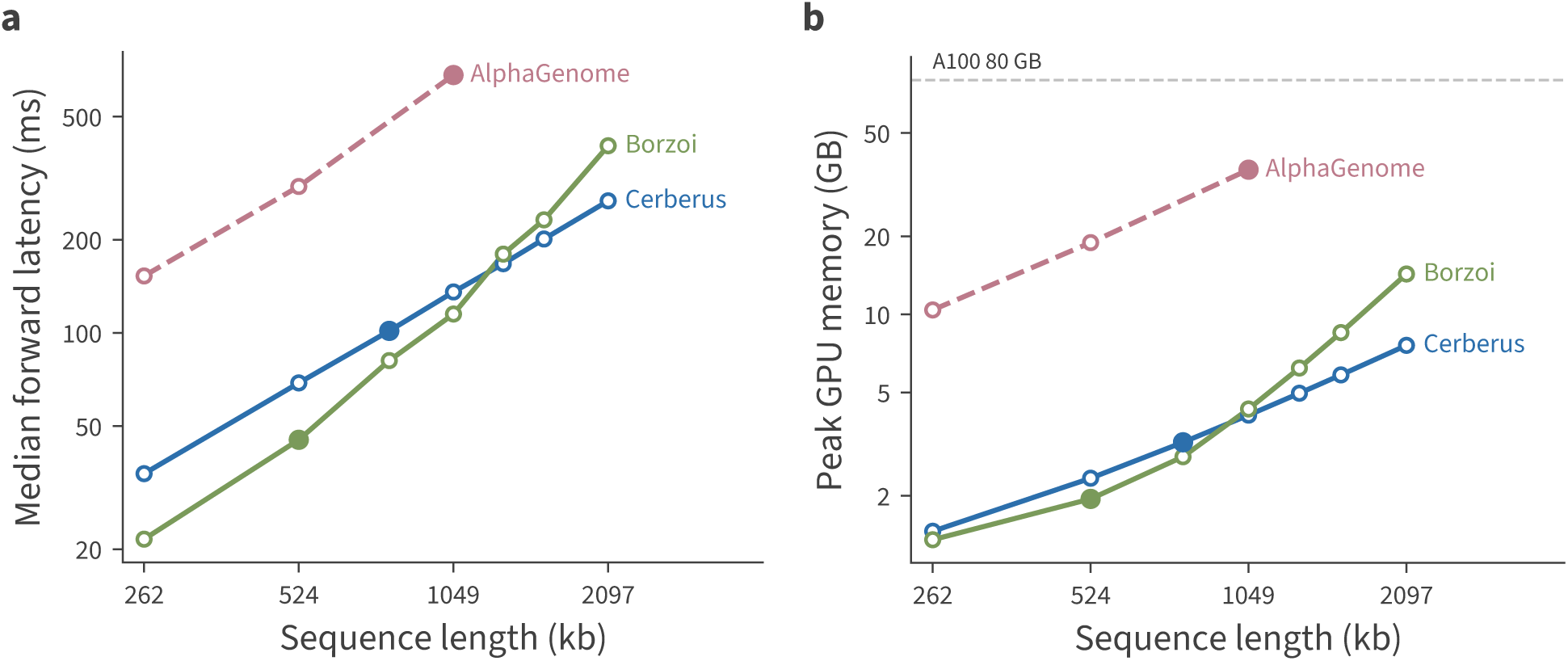
Inference cost against sequence length on an A100 80 GB. Cerberus and Borzoi were measured on one GPU (Google Cloud a2-ultragpu-1g) from 262 kb to 2.10 Mb; AlphaGenome values come from a published benchmark of its official JAX implementation on the same GPU model [25], since its license does not permit running it outside the API. All series follow one protocol: batch size 1, three warmup passes discarded, median of ten timed forward passes, variable-size prediction heads inside the timed path. Each series is plotted with its fastest implementation, torch.compile with bfloat16 autocast for Borzoi and Cerberus and JAX for AlphaGenome; the published benchmark times a compiled step throughout, and compilation beat eager execution in every cell of our sweep. Filled markers sit at each model’s training length, 786 kb for Cerberus, 524 kb for Borzoi and 1 Mb for AlphaGenome; open markers are other lengths, and the published AlphaGenome measurements stop at 1 Mb. **(a)** Median forward latency, scaling as L^0.98^ for Cerberus and L^1.40^ for Borzoi, as expected of a linear-time sequence mixer against attention. The curves cross between 1.05 and 1.31 Mb. **(b)** Peak GPU memory, from torch.cuda.max memory allocated, scaling as L^0.80^ and L^1.14^ and crossing earlier, between 786 kb and 1.05 Mb. The community PyTorch port of AlphaGenome (not plotted) is reported 4–13% slower on 10–13% more memory than the JAX implementation. Cerberus and Borzoi were run in one harness, so their comparison is controlled; AlphaGenome was measured by other authors and carries the added uncertainty of a separate harness on a separate GPU of the same model.

**Fig. S13.**
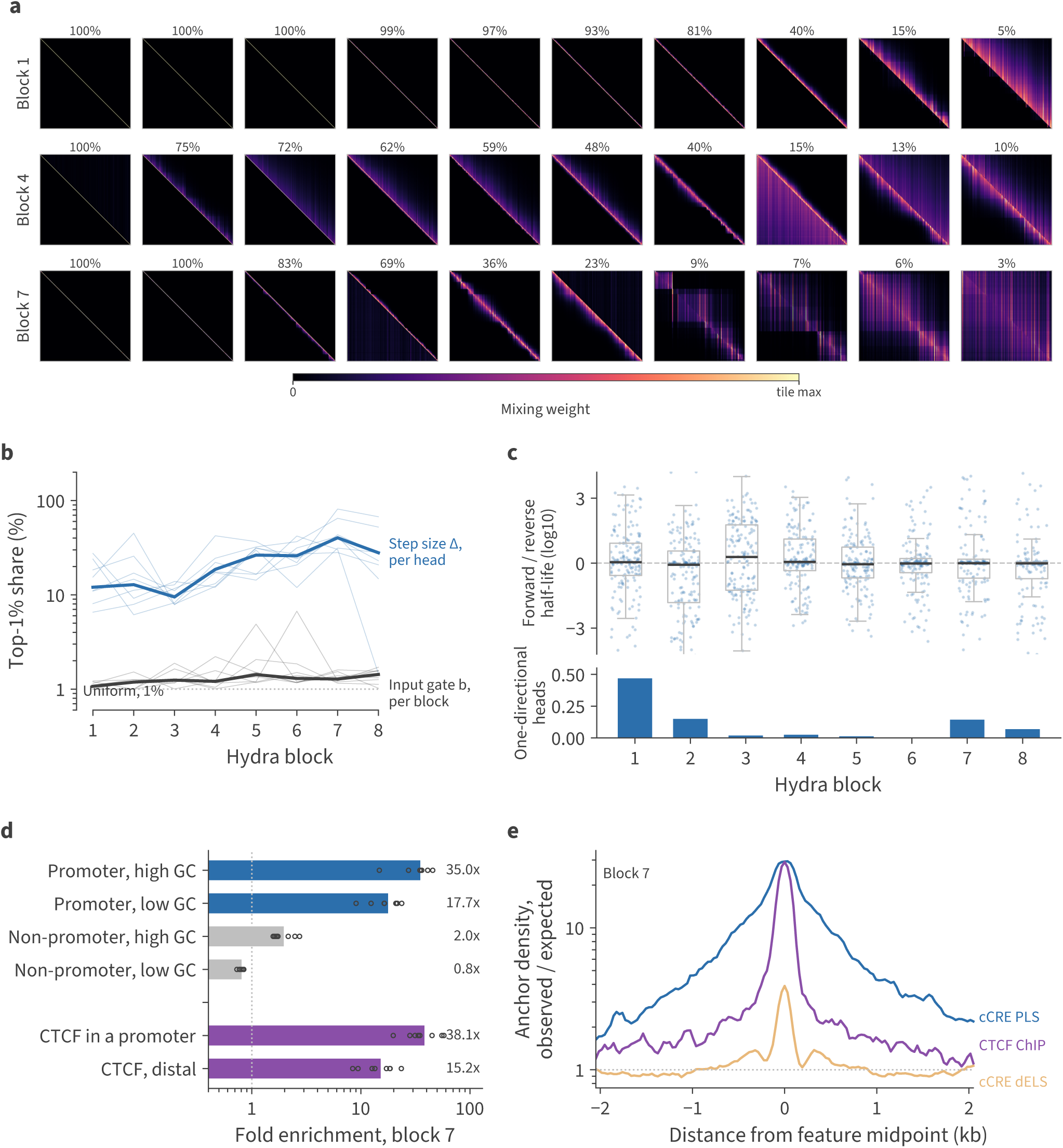
Hydra head geometry, directionality, and controls on the anchor enrichment. Panels cover the eight Cerberus replicates, each with 8 blocks of 20 heads, on 16 held-out 786 kb sequences. **(a)** Every other head of the first, fourth and seventh blocks in replicate 1, on the held-out window of Fig. 4d. Heads are ordered left to right by on-diagonal weight, the share of *|m_ij_|* within two cells of the diagonal, printed above each tile. The first block is almost all diagonal, the fourth banded, and the seventh mixes diagonal, banded and column-anchored heads. Each tile is scaled to its own 99.9th percentile, since the output projection absorbs the absolute scale of a head’s operator, so the bar below the grid runs from zero to each tile’s own maximum. **(b)** Share of each track carried by its top 1% of positions, by block, for the per-head step size Δ and the input gate b. Heavy lines are the median over all replicates and heads, thin lines the per-replicate medians. Because ngroups = d_state_ = 1, b is one channel shared by all 20 heads of a block, and it stays near the 1% a flat track would give at every depth. **(c)** Top, the log_10_ ratio of forward to reverse half-life within a head, one point per head with boxes spanning the interquartile range. Bottom, the fraction of one-directional heads, in which one pass’s mean absolute output is under 1% of the other’s. **(d)** Anchor enrichment in the seventh block after splitting each class on its confound. The seventh block is shown rather than the eighth because its step size concentrates more, in all 8 replicates, and its half-life is longer; the two blocks’ enrichment barely differs. Promoter-like cCREs and non-promoter bins are split at the median GC content of promoter-like cCREs; CTCF ChIP-seq peaks are split on overlap with a promoter-like cCRE. Bars are the mean over replicates with the individual replicates overlaid. **(e)** Anchor density in 32 bp bins against signed distance to the midpoint of the nearest feature, seventh block, observed over the density expected from the class’s rate in the scored windows.

**Fig. S14.**
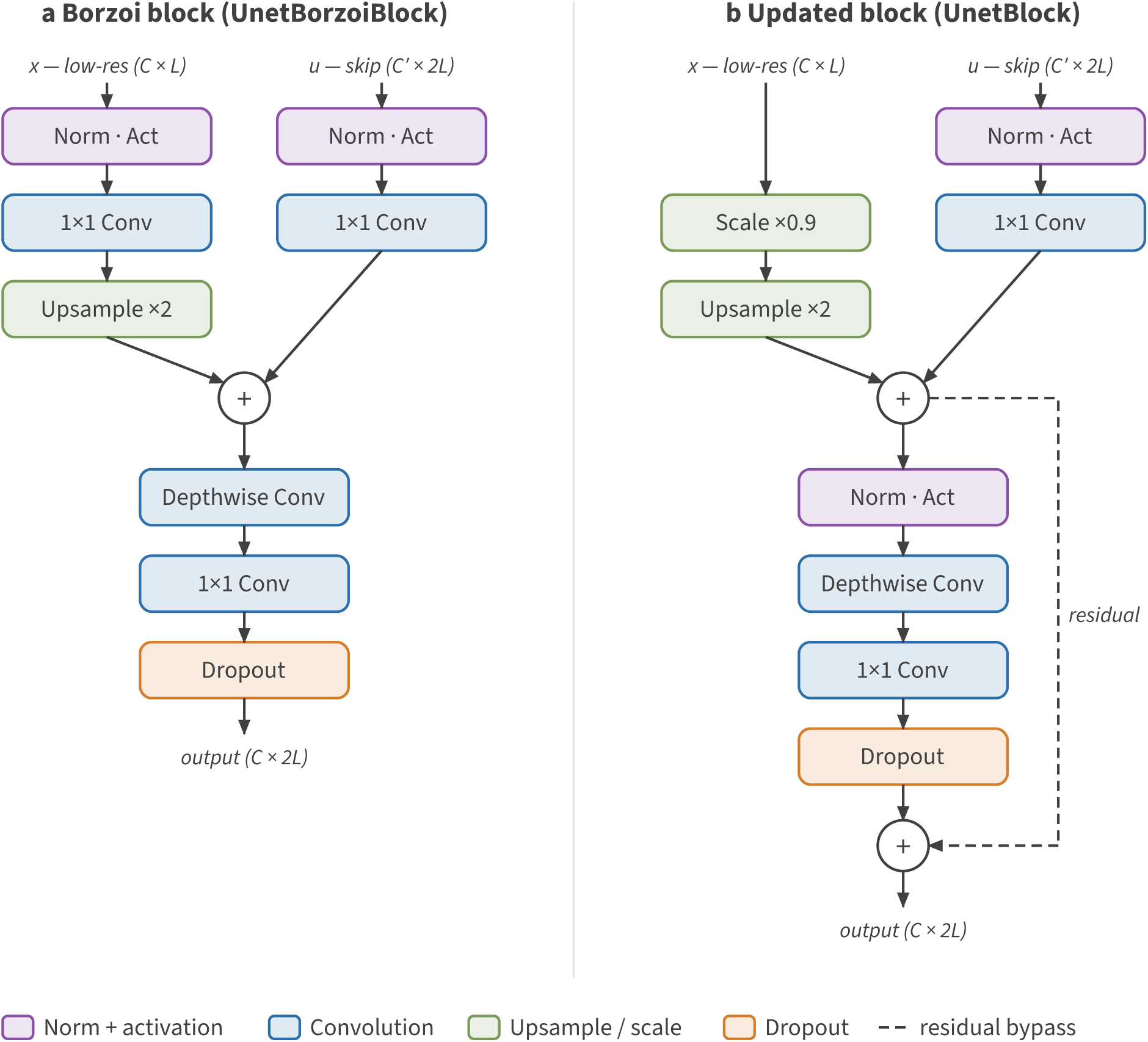
U-net decoder block. Borzoi versus updated upsampling U-net block. The new block scales the coarse path by a learnable factor instead of normalizing/convolving it, normalizes the merged sum rather than each branch, and adds a residual connection around the depthwise-separable refinement; the skip path is unchanged.

**Fig. S15.**
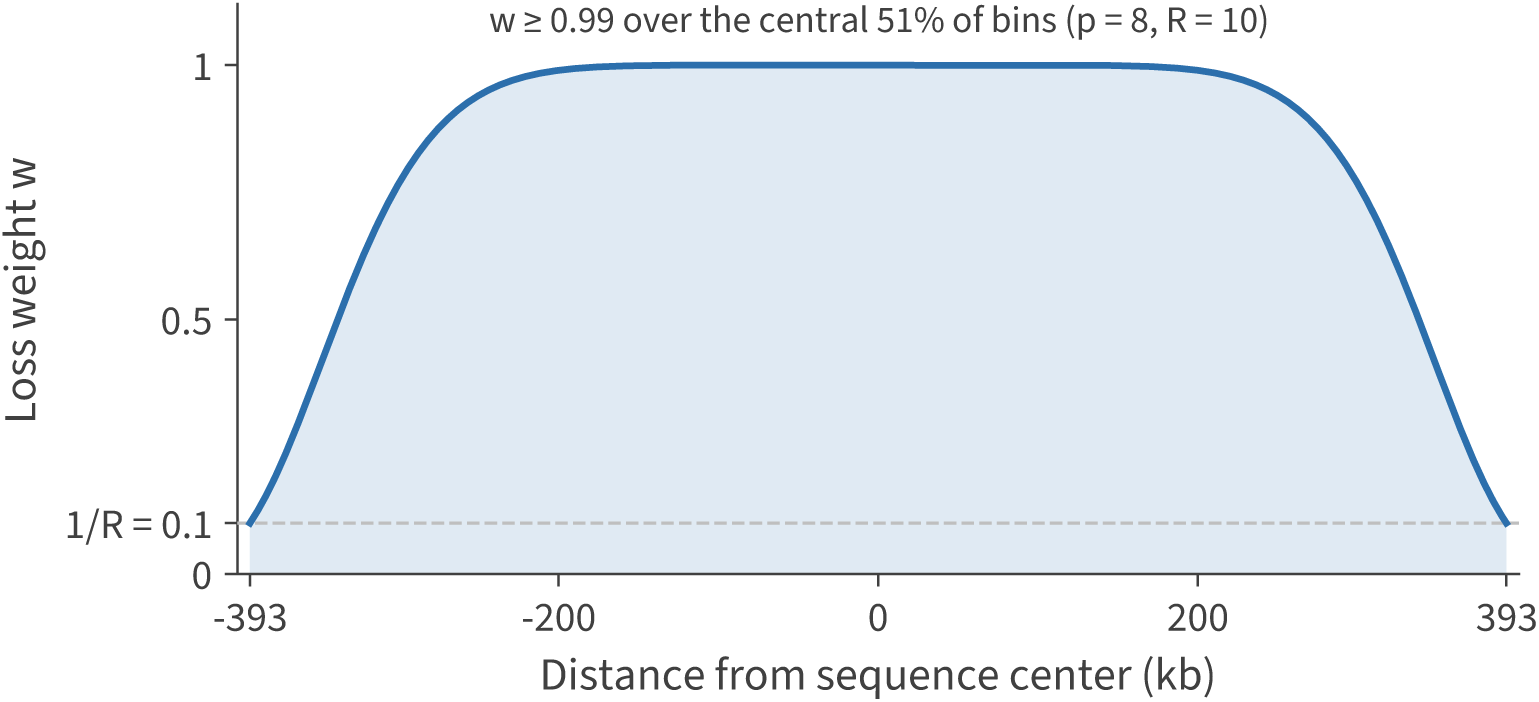
Spatial position-weighted loss. Each output bin’s contribution is scaled by a super-Gaussian window *w* = exp[*−*(*u*/σ)*^p^*], where *u* is the bin’s distance from the sequence center. With *p* = 8 and center-to-edge dynamic range R = 10 (the values used for all training runs), the weight holds a flat plateau at *≈* 1 across the well-supported interior and rolls off smoothly to 1/R = 0.1 (dashed line) at the edges, down-weighting boundary bins whose receptive fields extend beyond the input window.

## Notes

https://github.com/calico/baskerville

